# miR-9 mediated noise optimization of the her6 oscillator is needed for cell state progression in the Zebrafish hindbrain

**DOI:** 10.1101/608604

**Authors:** Ximena Soto, Veronica Biga, Jochen Kursawe, Robert Lea, Parnian Doostdar, Nancy Papalopulu

## Abstract

Ultradian oscillations of key transcription factors, such as members of the Hes family, are thought to be important in Neural Progenitor Cell (NPC) maintenance and miR-9 acts as a tuner of these oscillations in vitro. However, the existence and the role of such dynamic oscillatory expression in vivo is poorly understood. Here, we have generated a Zebrafish CRISPR knock-in Her6::venus fusion (Hes1 orthologue) to study endogenous dynamic gene expression in the embryonic hindbrain. We show that Her6 undergoes a transition from irregular, noisy, fluctuations to periodic oscillations as neurogenesis proceeds. In the absence of miR-9 input, noise in the Her6 oscillator increases and NPCs are unable to transit away from an intermediary state where they co-express progenitor and early differentiation markers. Thus, Her6 oscillations are facilitated by noise optimization mediated by miR-9 and this noise-tuning step is functionally important for cells to transition to differentiation.

## Introduction

Understanding how cells make cell state transitions from neural stem/progenitor cells to differentiated neurons is key to interpret most biological processes that underlie the development of the nervous system. Indeed, cell state transitions underlie the development of most biological systems, regeneration and cancer. Transcriptomic analysis of single cells is a very powerful method that has been used to understand cell state transitions as it can uncover thousands of genes that are up or down regulated in each state. Aided by sophisticated bioinformatic tools, transcriptomic analysis can also reveal the path taken by cells and the inferred temporal order (“pseudotime” as in e.g.^1,2^). However, while such transcriptomic analysis is powerful in elucidating a sequence of gene expression it is essentially a snapshot analysis and as such it does not reveal the fine-grained dynamics of gene expression that take place in a timeline of just a few hours.

Are such relatively short-scale dynamics important? Evidence is mounting that pulsatile or oscillatory expression that takes place in an ultradian scale carries information encoded in its characteristics, which can be decoded by downstream processes^3^. Although the power of pulsatile gene expression is recognized in many systems across biology, so far, examples in developmental biology are few. Leading developmental examples are the ultradian oscillations of Transcription factors (TFs) and signaling molecules in the context of somitogenesis and neurogenesis. Key examples include Hes genes and proneural TFs; e.g. Ascl, Olig and ngn, members of Notch signaling (e.g. delta; ^4–6^ reviewed in ^7^) and wnt signaling pathways ^8^. Wherever it has been tested by experimentation, it was clear that sustained versus pulsatile expression of such molecules has distinct outcomes for cell fate decisions ^9^.

For the development of the nervous system understanding the dynamics of Hes gene expression is particularly important because TFs of this family are important for neural progenitor maintenance and controlled differentiation ^10^.

Oscillatory dynamics can be revealed with live imaging using protein reporter fusions. The value of such protein-reporter fusions is that they are more likely to recapitulate the properties of the endogenous proteins, many of which are highly unstable. Indeed, instability of components (mRNA or protein) are an essential property in several biological oscillators ^11^. In neurogenesis, such protein fusions have been invaluable in characterizing the oscillatory dynamics of *hes* genes, proneural genes (Ascl, ngn and Olig2) as well as Dll (reviewed in ^7^). However, with few exceptions (e.g. ^6^), most of these studies have been performed in dissociated cells, cultured in 2D. Recent evidence in the mouse suggests that the 3D tissue environment can modify the oscillatory dynamics ^12^, therefore, it is essential to be able to study protein expression dynamics in vivo.

Zebrafish is ideal for such in vivo studies because of its superior suitability for live imaging of molecular and cellular events at several time scales. This has been exploited in the context of oscillations during somitogenesis, both at the population and single cell level ^13–15^. However, virtually nothing is known about the real-time dynamics of gene expression during Zebrafish neurogenesis, although previous studies based on fixed tissue snapshot analysis have implied a highly dynamic interaction of *her* genes, their targets and regulators ^16,17^. Specifically, *her* genes have been implicated in neural progenitor maintenance and in particular, in maintaining cells in an ambivalent progenitor state ^17^. These transitions are controlled by miR-9 ^16,17^ as is the case also in the mouse ^18^ mediated by controlling Hes dynamics ^19,20^. Thus, there is need to understand *her* gene expression dynamics in vivo and the potential role of miR-9 in tuning these dynamics.

Here, we use CRISPR/Cas9 technology to create the first fluorescent moiety knock-in Zebrafish to be used beyond proof of principle ^21^ for experimental purposes. We knocked-in Venus fluorescent protein in frame with Her6, a Hes1 homologue, and after thorough characterisation of the reporter, we used it to study the endogenous Her6 protein dynamics in Zebrafish hindbrain neurogenesis. We find that Her6 is initially expressed in neural progenitor cells in a fluctuating and noisy but not periodic manner. Oscillations with regular periodicity appear at the peak of neurogenesis and coincide with the onset of expression of miR-9 in the hindbrain, consistent with a previous reported input of miR-9 in regulating Hes1/Her6. To investigate the precise function of miR-9 in tuning Her6 dynamics, we use CRISPR/Cas9 to mutate the miR-9 binding site in the 3’UTR of *her6*. We report that preventing the influence of miR-9 on Her6 increases the amount of protein expression noise and decreases the quality of oscillations during neurogenesis. Surprisingly, under these conditions, cells also fail to down regulate Her6, to upregulate proneural genes and to differentiate; instead they accumulate in a transitory state which co-expresses progenitor and early neuronal markers. These findings allow for the first time the real-time imaging of protein expression dynamics at the ultradian scale and suggests that miR-9 is necessary to reduce noise in the Her6 oscillator. Taken together, we suggest that miR-9 mediated optimization of noise allows cells to produce regular Her6 oscillations and enables cell state transitions to differentiation to occur.

## Results

### A Her6::Venus knock-in protein fusion is a faithful reporter of endogenous Her6 expression and dynamics

In order to characterise the dynamics of cell state transitions, we aimed to identify the most suitable Zebrafish *her* gene for dynamic analysis of gene expression. There are two *hes1* related genes in Zebrafish; *her6* and *her9* ^22^. They are both expressed in the Zebrafish embryonic central nervous system (CNS; Fig. 1a.i-ii) mostly in a mutually exclusive pattern appearing as adjacent narrow “bands” of cells that span the dorso-ventral axis in the hindbrain (Fig 1a.iii). Both *her6* and *her9* harbor a miR-9 binding site in the 3’UTR but the *her6* site is a better quality-binding site (7A1-mer rather than 6-mer; Fig. 1b), therefore we decided to focus on Her6.

**Figure 1.**
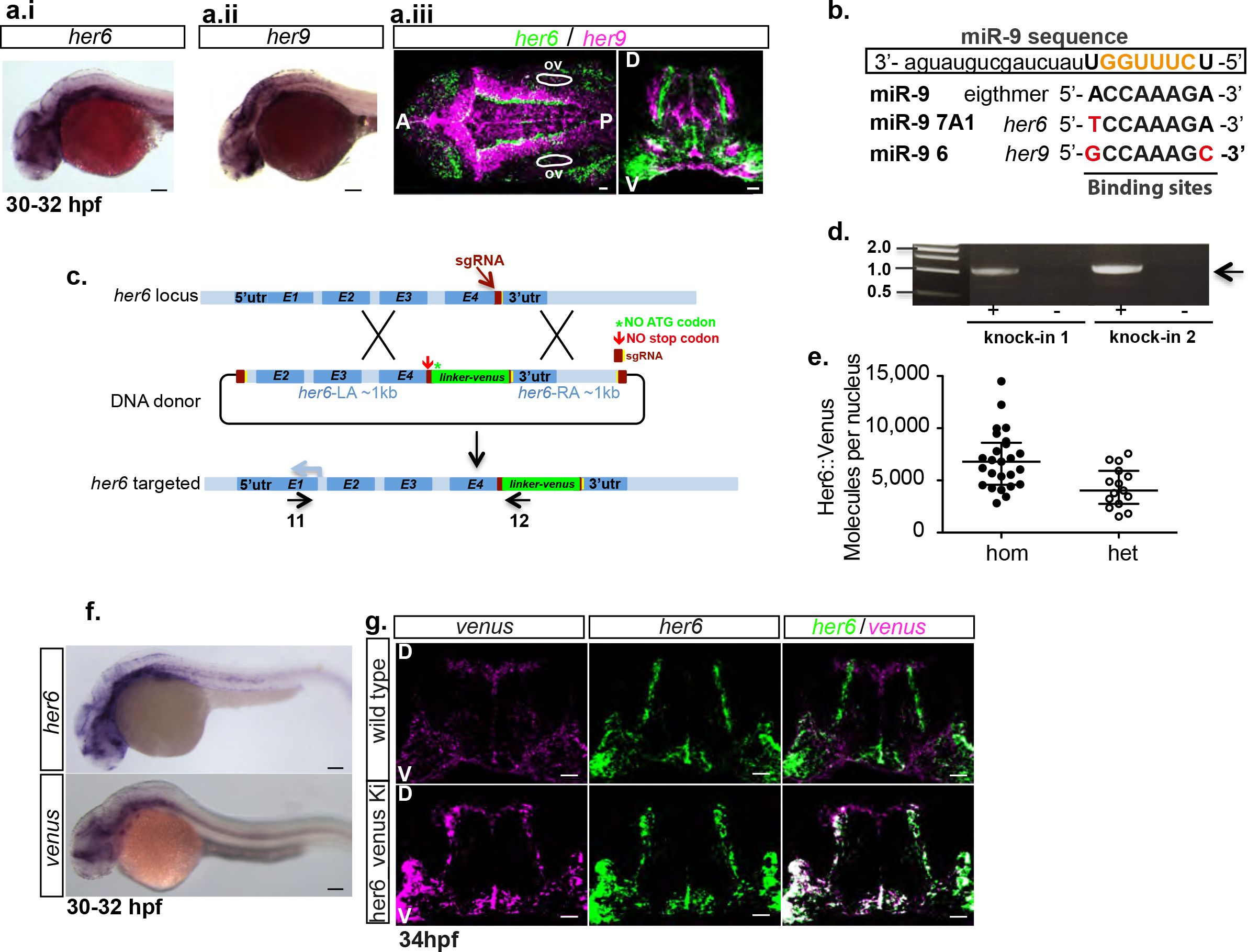
Her6::Venus knock-in generation and characterization. **(a)** Chromogenic whole mount in situ hybridization (WMISH) for *her6* **(a.i)** and *her9* **(a.ii)**; longitudinal view, scale bar represents 100μm. **(a.iii)** Double fluorescent WMISH to detect *her6* (green) and *her9* (magenta); coronal view (left panel) and transversal section (right panel), scale bar 20μm; 30-32 hpf development stage. **(b)** *her6* and *her9* microRNA-9 binding site sequence compared to the complementary eigthmer sequence of MiR-9, highlighted in red are the mutations occurring in each *her* gene. The boxed sequence correspond to the miR-9 sequence, in bold and capital letter is the binding sequence. **(c)** Schematic showing the strategic approach to generate the Her6::Venus knock-in; annotations indicate LA: left arm, RA: right arm. **(d)** Amplicon generated by using primers 11 and 12 using schematic in **(c)**, obtained only when Venus is inserted at C-terminus of *her6* gene; (+): Her6::Venus knock-in embryo, (−): wildtype sibling embryo. Knock-in 1 and 2 are two different founder animals. **(e)** FCS absolute quantitation of Her6::Venus protein abundance observed in the hindbrain, rhombomere 6, of homozygous versus heterozygous embryos at 34±1hpf; bars represent median and interquartile range of (hom: 25cells) and (het: 15cells) collected from one embryo per condition. **(f)** Chromogenic WMISH for *her6* in wild type embryos (top panel), and *venus* in Her6::Venus knock-in embryos (bottom panel); development stage 30-32 hpf; longitudinal view, anterior to the left, scale bar 100μm. **(g)** Transverse sections of double F-WMISH for *her6* (green) and *venus* (magenta) observed in wild type embryo (top panel) and Her6::Venus knock-in embryo (bottom panel) at 34hpf showing that *her6* and *venus* expression coincide in the hindbrain of the knock-in reporter animals; scale bar 20μm. Image annotations indicate A: anterior, P: posterior, D: dorsal, V: ventral, ov: otic vesicle.

To generate a reporter that would be suitable for live imaging, we devised a CRISPR/Cas9 knock-in strategy that would preserve the properties of the endogenous protein (suppl. Fig. S1a). This involved an in frame fusion of the fluorescent moiety, *venus*, to the C-terminus of the endogenous *her6*, placed before the 3’UTR (Fig. 1c and Online Methods for details on DNA donor design and guideRNA selection), anticipating that destabilization sequences within the Her6 protein would similarly destabilize the fusion protein. While the *venus* stop codon was preserved, the stop codon of the endogenous *her6* and the start codon of the *venus* were removed and a small linker was used to allow both proteins to independently fold (Fig. 1c).

The frequency of repaired DNA with the insertion of *venus* in F0 fish was 30% with a decrease to 3.3% (2/60) from those F0 adults carrying germ line transmission, identified by the PCR fragment of the expected size (Fig. 1d, suppl. Fig. S1b.).

Sequencing of F1 embryos throughout the endogenous-donor DNA fusion was used to ensure that the reading frame was correctly maintained (suppl Fig. S1c. ii). The F1 embryos were selected for the presence of the reporter by fin clipping at 3dpf and genotyped by qPCR making use of the derivative of the melting curve (suppl. Fig. S1d.i-iii).The positive fish were confirmed by fin clipping at 8wpf and visualization of the right size amplicon (suppl. Fig. S1a, suppl. Fig. S1d.iv and Online Methods for strategy to obtain F1 generation).

To ensure that the reporter recapitulated accurately the expression of Her6, a number of tests were carried out. The protein molecule number was identified in single hindbrain cells by Fluorescence Correlation Spectroscopy (FCS) in homozygous and heterozygous embryos and the ratio was found to be 1.8, indicating that additional integrations into the genome are unlikely (Fig. 1e and suppl. Fig. S1c.iv). The mean number of molecules in the homozygous fish was 7,000 protein molecules per nucleus, at stage 30-34hpf, which indicates that Her6 protein is a low abundance protein (Fig. 1e), similar to the mouse Hes1 in neural progenitor cells ^19,23^. The expression of Venus was compared to the expression of endogenous Her6 by chromogenic and fluorescent whole mount in situ hybridization (WM-FISH) and sections through the hindbrain. Neither ectopic nor any region of missing expression were identified (Fig. 1f,g). There was no significant change in the somite number between control, heterozygous or homozygous Her6::Venus embryos at 72hpf (suppl. Fig. S1c. iii), suggesting that the knock-in reporter does not interfere with normal development. Finally, there was no significant change in the protein half-life of HA-Her6 and Her6::Venus, both of which were very unstable (average half-life 12 and 11min, respectively; suppl Figure S1e). These findings confirm that the Her6::Venus fusion protein is a faithful reporter for visualising endogenous Her6 dynamic expression.

### Her6::Venus protein expression is dynamic during development, reflecting active hindbrain neurogenesis

Little is known about the dynamics of Her6 gene expression during neurogenesis since only fixed samples analysed by in situ hybridization at stages earlier than 24hpf have been described before ^24^.Thus, we first focused on characterising the Her6 protein expression in the hindbrain during the period of development when neurogenesis takes place, i.e. between 20 and 48hpf ^25^.

Still images of the lateral view from the Her6::Venus knock-in brain reveal that Her6 is expressed in the hindbrain rhombomeres (r1-r6) with an overall anterior-posterior gradient mode of intensity (Fig. 2a.i and Movie S1). A temporal gradient is also observed since expression starts in r6/r7 at 20hpf and spreads anteriorly to r1, r2, r3, r4 and r5, maintaining a high level of expression in r6 (Fig. 2a.i).

**Figure 2.**
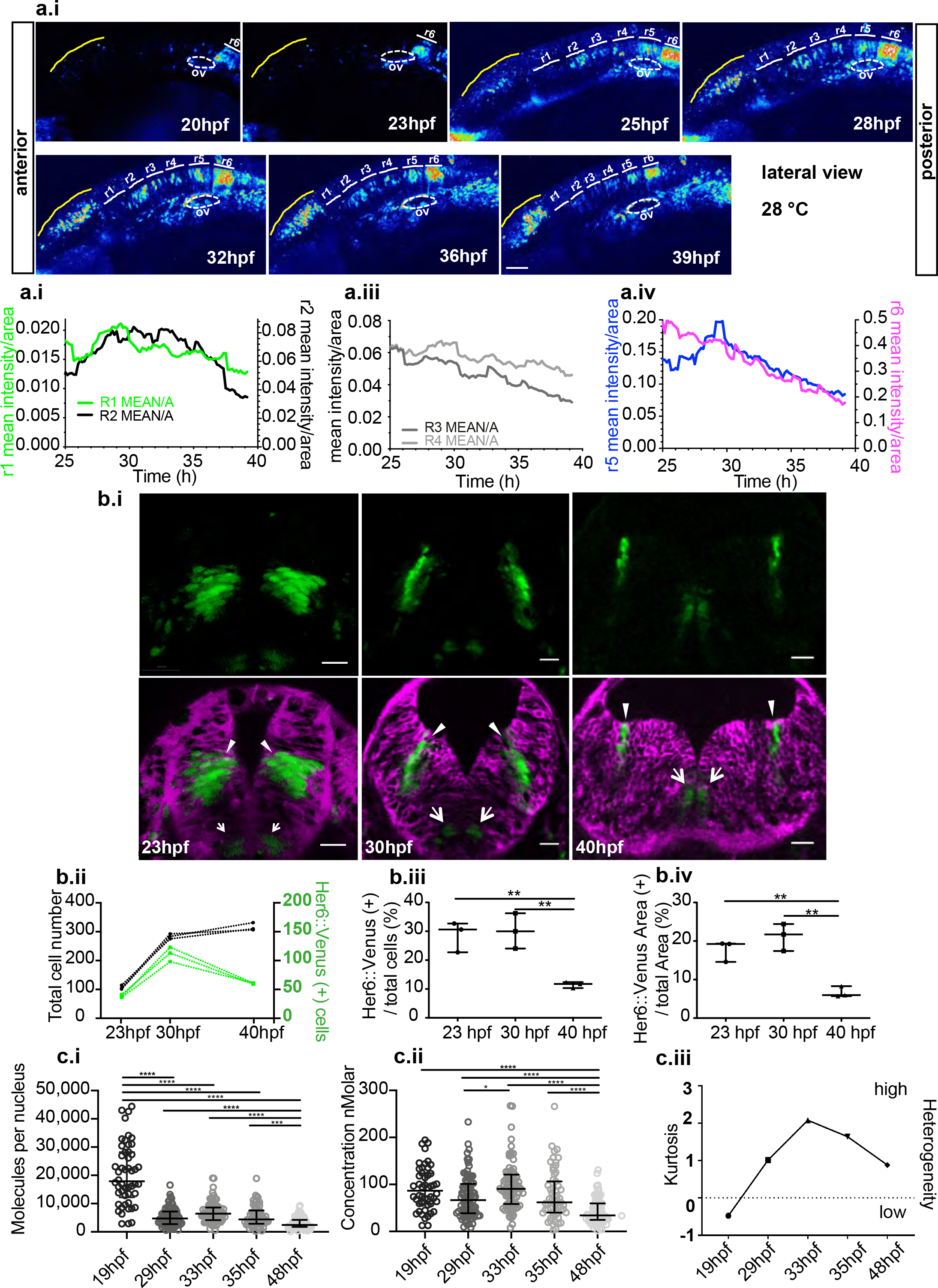
Her6::Venus protein expression is dynamic during development, reflecting active hindbrain neurogenesis. **(a.i)** Time series example of Her6::Venus expression during development, in the midbrain and hindbrain; longitudinal view; scale bar 50μm; also included in Movie S1. **(a.ii-iii)** Intensity mean of Her6::Venus per rhombomere area over development with high expression in r5 and r6 shown separately. **(b.i)** Transverse view snapshots in hindbrain of Her6::Venus embryos showing r6 over time; Her6::Venus protein has two expression domains: a ventral domain (arrows) and a more dorsal lateral domain (arrowheads); the caax-mRFP was used as membrane marker (magenta); scale bars 20μm; images at 30-40hpf are maximum projection of 4 z-stack from Movie S2 (16 z-stacks). **(b.ii)** Quantification of Her6::Venus(+) cell number (green) compared to total cell number (black lines) over development. **(b.iii-iv)** Proportional changes in Her6::Venus(+) cell numbers **(b.iii)** and Her6::Venus(+) area **(b.iv)** during development indicate a significant reduction of her6 expression domain which coincides with a reduction in number of expressing cells; bars indicate median and interquartile range of counts collected from 3 z-stacks per embryo and one embryo per condition; statistical tests indicate one-way ANOVA with Bonferroni multiple comparison with significance **(p<0.01). **(c.i-ii)** Nuclear abundance **(c.i)** and nuclear concentration **(c.ii)** of Her6::Venus protein in homozygous embryos at different stages during development measured by fluorescence correlation spectroscopy (FCS) shows the absolute molecule number per cell and indicates downregulation during development; bars indicate median and interquartile range of (19hpf: 6 embryos, 50 cells), (29hpf: 6 embryos, 86 cells), (33hpf: 5 embryos, 76 cells), (35hpf: 6 embryos, 59 cells) and (48hpf: 6 embryos, 89 cells); statistical tests indicate Kruskal-Wallis with Dunn’s multiple comparison with significance * (p<0.05), ***(p<0.001) and ****(p<0.0001). **(c.iii)** Quantification of heterogeneity using kurtosis from concentration data in **(c.ii)** indicating maximal heterogeneity at 33hpf; null kurtosis corresponds to a normal distribution.

Individual rhomobomeres show variable levels of Her6, lower in r1/r2/r3/r4 compared to r5/r6 (Fig. 2a.ii-iii vs a.iv). Each rhombomere has a distinct profile of Her6 dynamic expression over time (Movie S1). Between 25hpf to 31hpf expression in r1/r2/r5 shows upregulation (Fig. 2a.ii and iv) meanwhile r3/r4/r6 have fluctuating but steady declining levels (Fig. 2a.iii and iv). Importantly, all rhombomeres exhibit protein down regulation over time and this conincides with the inflection point when neurogenesis starts increasing exponentially in the hindbrain ^25^. The Her6 expression profile is higher in r5/r6 with r6 having a constant slow decline over time from 25hpf, reflecting overall the expected Her6 downregulation as cells differentiate and suggesting a prolonged neurogenesis. We selected rhombomere 6 for further dynamic analysis because, in spite of the overall steady decline, Her6 is highly expressed in r6 and the otocyst can be used as a clear anatomical landmark.

Transversal views of the Her6::Venus expression in the hindbrain (r6) showed expression in two restricted domains, each starting close to the ventricular zone and extending further out towards the basal surface (Fig. 2b and Movie S2). A ventral domain potentially contributes to motor neuron circuits (Fig. 2b.i arrows) ^26^, while the more dorsally located domain, halfway along the D-V axis, is likely to encompass interneuron progenitors ^26^(Fig. 2b.i arrowheads). This dorsal domain was the subject of subsequent investigation. The number of Her6 expressing cells within this domain initially increases in absolute numbers but not the percentage of expressing cells (Fig. 2b.ii-iii 23hpf vs 30hpf); meanwhile, in the later phases of neurogenesis we noted both an absolute and proportional decrease in Her6 expressing cells (Fig. 2b.ii-iii 30hpf versus 42hpf) perhaps reflecting a switch from symmetric (i.e. proliferative) to asymmetric (i.e. neuron-generating) divisions of Her6-positive NPCs. There is no reduction of the Her6 domain area between 23 and 30hpf despite the apparent morphogenetic movements, while reduction between 30 and 42hpf (Fig. 2b.iv 30hpf vs 42hpf) is due to actual reduction of NPC numbers (Fig. 2b.i-iii 30hpf vs 42hpf). When we quantified the absolute protein molecule number per nucleus by FCS, we observed the highest abundance at 19hpf (Fig. 2c.i); this is a consequence of a larger nuclear volume observed at this stage (Fig. 2b.i, 23hpf versus 30hpf) and no significant difference in concentration was seen (Fig. 2c.ii, 19hpf versus 29hpf). We also observed a wide heterogeneity in the Her6 concentration at the single cell level that had its peak around 33hpf (Fig. 2c. ii-iii; Online Methods).

In summary, we observe a dynamic Her6 expression at a population level with a declining trend on expression, reflecting NPC differentiation. This encompasses high gene expression heterogeneity that could be due to dynamics at single cell level over time and is studied next.

### Her6::Venus dynamic expression with single cell resolution in the Hindbrain shows oscillations and noise

To characterise the dynamic expression of Her6 in normal hindbrain development, we used continuous live imaging over 12hr of Her6::Venus homozygous reporter embryos. We injected mKeima-H2B and caax-mRPF mRNAs to serve as nuclear and membrane landmarks, respectively, to facilitate segmentation of individual cells (Fig. 3a, suppl. Fig. S2a, Online Methods-Image processing).

**Figure 3.**
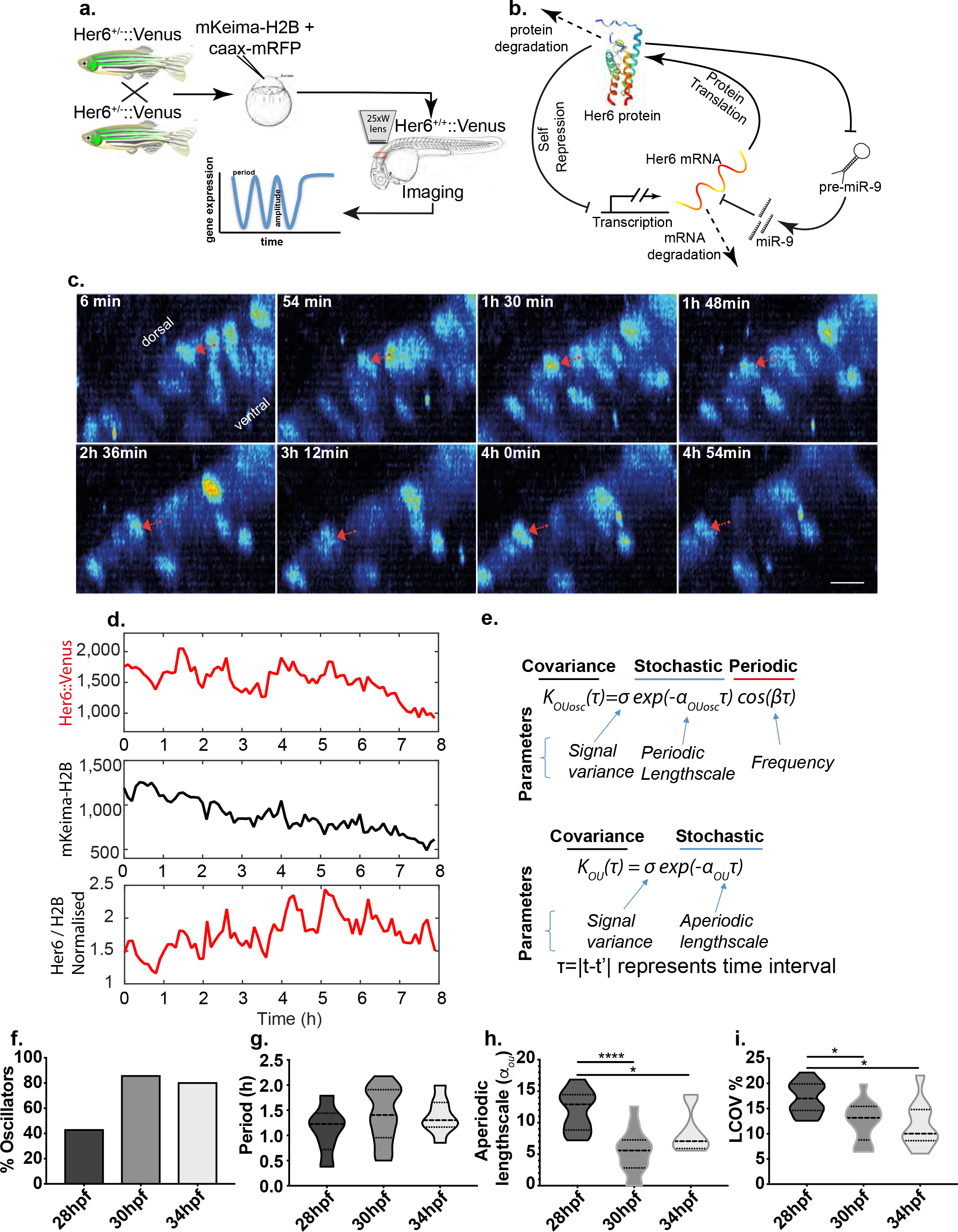
Changes in dynamics of Her6::Venus in single neural progenitor cells during development. **(a)** Diagram of experimental approach to image Her6::Venus dynamic expression at single cell resolution. **(b)** Schematic representation of genetic auto-repression network of Her6 including microRNA-9 regulation. **(c)** Representative time lapse images of a single neural progenitor cell (red arrow) tracked in 3D data starting from 34hpf, showing intensity fluctuations over time; scale bar 10 μm; images corresponding to Movie S3. **(d)** Single cell time series of Her6::Venus, mKeima-H2B and Her6::Venus signal normalized by mKeima-H2B corresponding to the cell in **(c)**. **(e)** Covariance models and parameters used to characterize oscillatory (*K*_*OUosc*_) and non-oscillatory (*K*_*OU*_) single cell expression. **(f-i)** Quantification of single cell dynamics at different stages in normal development including: **(i)** proportion of oscillatory cells, **(g)** period of oscillators estimated using *K*_*OUosc*_, **(h)** Noise measured by aperiodic lengthscale (α_OU_) of the *K*_*OU*_, and **(i)** changes in local amplitude around the mean by coefficient of variation (LCOV); bars indicate median with interquartile ranges of 10-14 cells per embryo per condition; statistical tests represent Mann-Whitney two-tailed with significance * (p<0.05), ****(p<0.0001).

Her6, like its mammalian counterpart Hes1, may generate oscillatory expression in the ultradian scale (i.e. with periodicity of a few hours) due to molecular auto-repression of transcription, coupled with instability of the her6 mRNA and Her6 protein, and tuned by miR-9 (Fig. 3b) ^20,27^. Semi-automated tracking of Her6::Venus expressing cells produced Venus and mKeima intensity traces over time.

Analysis of the single cell time series of Her6::Venus showed fluctuations in intensity of expression (Fig. 3c, red arrow and Movie S3) which persisted when corrected for variability in the mKeima-H2B signal (Fig. 3d) and in combination with subtraction of long-term trend (detrended data-suppl Fig. S3a) we investigated the presence of ultradian periodicity in further analysis.

We interrogated the ability of progenitors to oscillate in Her6 levels over time using a statistical method previously developed to detect periodicity in Luciferase time series ^28^ and subsequently improved for noisy fluorescent data in mouse tissue ^12^. Our method uses sophisticated computational techniques to infer parameters of two Ornstein-Uhlenbeck (OU) covariance models *K*_*OUosc*_ and *K*_*OU*_ which are characteristic of periodic and aperiodic dynamics respectively (Fig. 3e). A strength of our approach is that we can classify cells into oscillatory and non-oscillatory with statistical significance (Online Methods-Periodicity analysis). Our covariance models include a lengthscale term that describes the rate of decay in correlation between subsequent peaks over time, referred to as periodic lengthscale *α*_*OUosc*_, and aperiodic lengthscale *α*_*OU*_. In addition to lengthscale, the periodic model also includes a cos wave term and this is characterised by frequency *β* and linked to period, *P* = 2*π*/*β*. Both models also account for the variance of the data which we analyse separately, hence, here was set to σ=1.

We used the stochastic *K*_*OUosc*_ covariance model to characterize Her6 oscillations at multiple embryonic stages (examples in Suppl. Fig. S3a). Our analysis showed that the proportion of cells that are classified as oscillatory greatly increases during development (40% at 28hpf, versus approx. 80% at 30 - 34hpf) (Fig. 3f) however there was no significant change in the periodicity of oscillators which had a median period of 1.2-1.4 h in all stages examined (Fig. 3g). Given that a large proportion of early stage progenitors were non-oscillatory (examples in Suppl. Fig. S3b), we then used the aperiodic covariance model, *K*_*OU*_ (Fig. 3e) to further investigate dynamics irrespective of ability to oscillate. Interestingly, fluctuations in Her6 expression in early progenitors were characterized by an increased aperiodic lengthscale which represents an increased rate of decay in signal auto-correlation over time (Fig. 3h). One can consider the rate of loss in correlation as an expression of “noise” and we conclude that early progenitors are noisier in their gene expression dynamics compared to later stage progenitors. We also analysed the local coefficient of variation (LCOV) denoting variability of signal around the mean; this is often taken as a measure of gene expression noise ^29^. The analysis of LCOV showed that early progenitors have higher gene expression variability than late progenitors (Fig. 3i 28hpf vs 30-34hpf).

Taken together, our findings suggest that the increase in neuronal differentiation observed during development of the hindbrain ^25^ is characterised by an increase in the number of cells that show oscillatory Her6 expression and an overall decline in the amount of protein expression noise, measured as an decreased rate (lengthscale) of correlation decay as well as decreased coefficient of variation.

### Change in Her6 dynamics by mutating the miR-9 binding site

The conversion from noisy to oscillatory expression as neurogenesis progresses implies a functional role for oscillatory Her6 gene expression. In Zebrafish, miR-9 has been found to target Her6, amplifying an ambivalent progenitor state that helps to fine-tune neurogenesis ^17^, however the effect on Her6 dynamics is not known. Previous work in 2D cultured mammalian cells suggested that miR-9 can tune the oscillatory expression of Hes1^18,27^, a finding that was further supported by computational modeling^19,20^. Therefore, we sought to make changes that will interfere or modify oscillatory expression of Her6 by changing the interaction with miR-9. In the Zebrafish hindbrain, miR-9 expression appears at 30-31hpf and continues to increase at least until 48hpf (Suppl. Fig S4a). The expression of miR-9 and Her6 spatially overlap, although the expression of miR-9 is wider (Fig. 4a), reflecting the existence of other targets ^30^. Based on these findings, we removed the influence of miR-9 on Her6 dynamics by mutating the miR-9 binding site in the *her6::venus 3’UTR*, to produce MicroRNA Binding Site mutant embryos (MBSm embryos, Fig. 4b). Experimentally, this was achieved by injecting sgRNA targeting the miR-9 binding site together with Cas9nls protein in one cell stage Her6::Venus Zebrafish embryos (supplem. Fig. S4b). Embryos were injected with minimal amounts of MBS sgRNA (F0) to not have overt phenotype at the macroscopic level during the experimental period (24hpf-52hpf), thus, minimizing the chances of non-specific toxicity.

**Figure 4.**
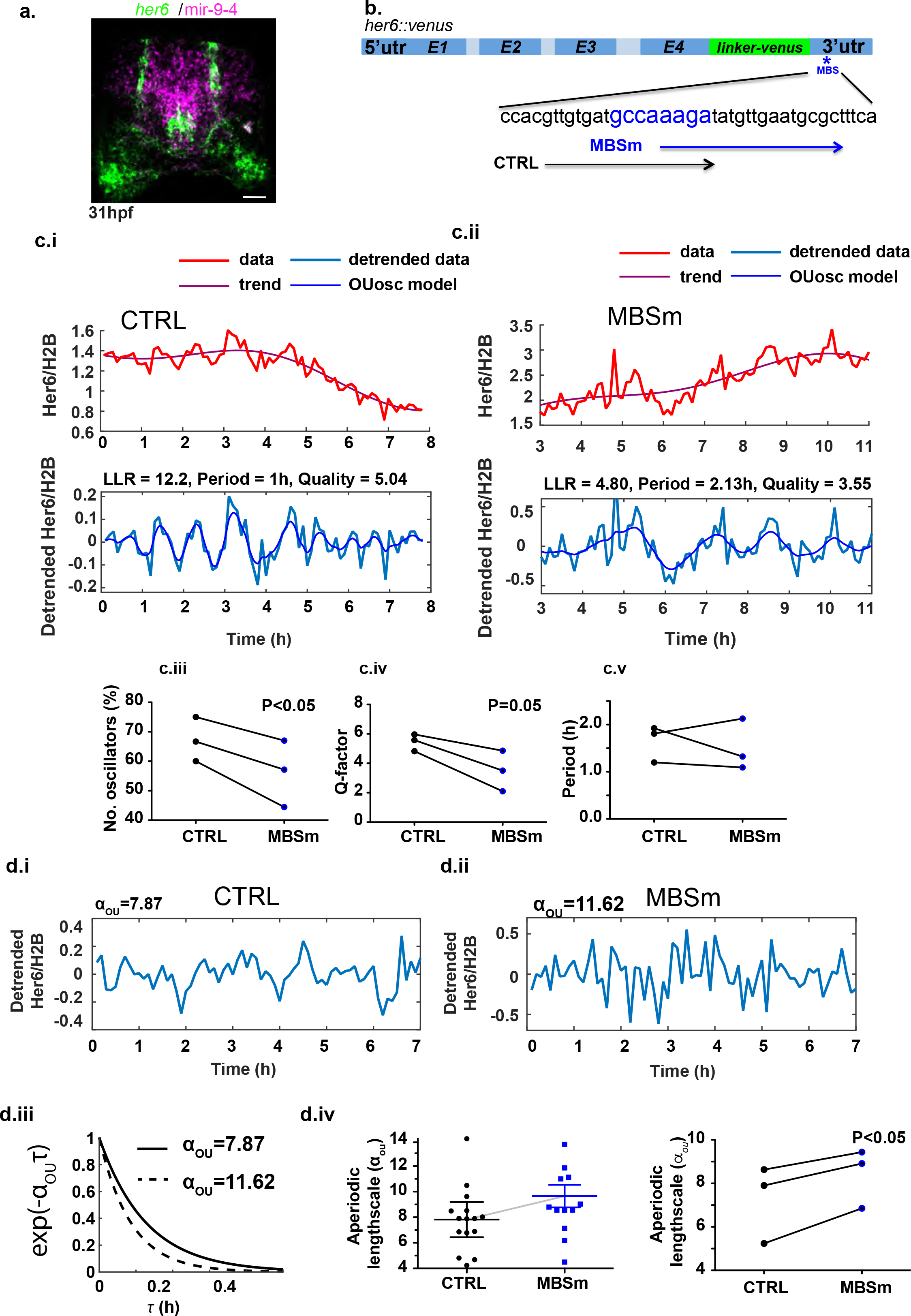
A mutation of the mir-9 binding site affects dynamics at single cell level. **(a)** Transverse section of double fluorescent whole mount in situ hybridization (F-WMISH) for *her6* (green) and mir-9-4 (magenta) imaged in fixed embryo at 31hpf; scale bar 30 μm. **(b)** Schematic representing the mir-9 binding site (MBS) of *her6::venus* mutated by CRISPR-Cas9nls protein; MBSm refers to specific sgRNA to mutate the MBS as opposed to sgRNA that does not produce mutation (control-CTRL). **(c.i-ii)** Representative examples of single cell oscillators observed in CTRL and MBSm embryos imaged from 34hpf onwards; time series represent Her6::Venus expression relative to mKeima-H2B (top panel) and detrended relative signal (bottom panel); parameters reported correspond to *K*_*OUosc*_. (**c.iii-v**) Pairwise analysis of oscillatory dynamics of control versus MBSm progenitor cells observed from 34hpf compared as median per experiment including: (**c.iii**) proportion of oscillators; (**c.iv**) Q-factor denoting quality of oscillations and **(c.v)** period of oscillators. **(d.i-ii)** Representative examples of detrended Her6::Venus relative to mKeima-H2B in non-oscillatory single cells observed in control and MBSm embryos imaged from 34hpf onwards. **(d.iii)** Quantification of noise by aperiodic lengthscale in one control versus one MBSm embryo with 12-14 cells per condition (top panel) and pairwise comparison of aperiodic lengthscale (bottom panel) shown as median per experiment; statistical tests indicate paired Student t-test with two-tail significance.

Co-injection of membrane bound mRFP (caax-mRFP) mRNA helped identify the injected embryos (Online Methods-microinjection and genotyping). High efficiency of mutagenesis of the miR-9 binding site (MBSm) (84% of injected embryos, n=7) meant that the phenotypic and dynamic analysis was possible in F0 embryos (suppl. Fig. S4b-c). Given that in uninjected embryos, heterogeneity of Her6 expression peaks around 34hpf (Fig. 2c.iii), and this correlates with a high propensity of progenitors to be oscillatory (Fig. 3f), we focused on the effects of MBSm at this stage of development.

Using our statistical framework (Online Methods-Periodicity analysis), we performed a single cell analysis of dynamics in traces observed in paired MBSm versus CTRL (control; inactive sgRNA injected) Her6::Venus embryos starting from 34hpf and imaged for approximately 12hrs (Fig. 4c.i-ii). We first quantified the propensity of Her6::Venus expressing cells to oscillate by using the periodic covariance model *K*_*OUosc*_ (Fig. 3e) and observed that in cells from MBSm embryos Her6 expression is less frequently oscillatory compared to control (Fig. 4c.iii). This is also associated with a persistent drop in the quality of oscillations in MBSm cells (Fig. 4c.iv and examples in Fig. 4c.i vs 4c.ii) quantified by a decrease in the Q-factor, which represents the ratio between frequency and periodic lengthscale. High values of Q-factor indicate similarity to a deterministic oscillator, and decreasing Q-factor values indicate irregularities in peak-to-peak variability and amplitude over time ^31, Phillips, 2016 #117^. The mean period of oscillators was not statistically significant in a paired test (Fig. 4c.v) indicating that both control and MBSm cells exhibit equivalent periods between 1h to 2h.

Since MBSm progenitors were more frequently classified as non-oscillatory compared to their non-mutated counterpart (Fig. 4c.iii), we then used the aperiodic covariance model, *K*_*OU*_ to describe the ‘speed’ of aperiodic fluctuations in the data by inferring the rate of correlation decay denoted by aperiodic lengthscale, *α*_*OU*_ (example non-oscillators shown in Fig. 4d.i-ii). We observed an increased rate of correlation decay over time in MBSm compared to control, indicating noisier Her6 expression in the absence of miR-9 regulation (Fig. 4d.iii-iv). Taken together, our findings are consistent with the miR-9 binding site mutation leading to a reduced quality and propensity of neural progenitors to oscillate and increased noise at the single cell level.

Having characterised dynamics in CTRL and MBSm progenitors we now turned our attention to the high frequency contribution to Her6 dynamics using power spectrum analysis. Outside the high frequency region, the power spectrum confirmed the presence of oscillations in both CTRL and MBSm embryos (Fig. S4d), with lower and fewer ultradian peaks in MBSm cells compared to CTRL cells (Fig. S4e) and consistent with observations in (Fig. 4c.iii-v). Within the high-frequency region, we observed an increased contribution of high frequency noise to variability of Her6::Venus dynamic expression in MBSm cells compared to control conditions (Figure S4f). Thus, both the analysis of aperiodic lengthscale (Fig. 4d.i-iv) and high frequency spectral characteristics indicate that gene expression is “noisier” in the absence of miR-9 input through its binding site on the *Her6* 3’UTR.

### Changes in fluctuation dynamics can affect downstream target expression

To understand how these observed changes in Her6 dynamic expression may affect cell state transitions, we sought to identify how higher levels of noise in the form of increased aperiodic lengthscales may affect the expression of downstream targets. We assumed that being a transcriptional repressor, Her6 (depicted as input Y in Fig. 5) represses the expression of a downstream target X, such as a pro-neural gene, which would be turned on when Y is downregulated. This minimal network-motif is highly sensitive to the aperiodic lengthscale in the dynamics of its repressing gene and assumes only one further gene regulatory interaction, namely that gene X may self-activate (Figure 5a). We computationally simulated this network motif using an ordinary differential equation, which allowed us to predict how downstream target expression of X may be affected by the change of upstream dynamics Y (Online Methods-mathematical modeling). We generated different hypothetical upstream dynamics by sampling traces from an OU Gaussian process characterized by the *K*_*OU*_ covariance function. This allowed us to directly vary the aperiodic lengthscale *α*_*OU*_ of the gene Y.

**Figure 5.**
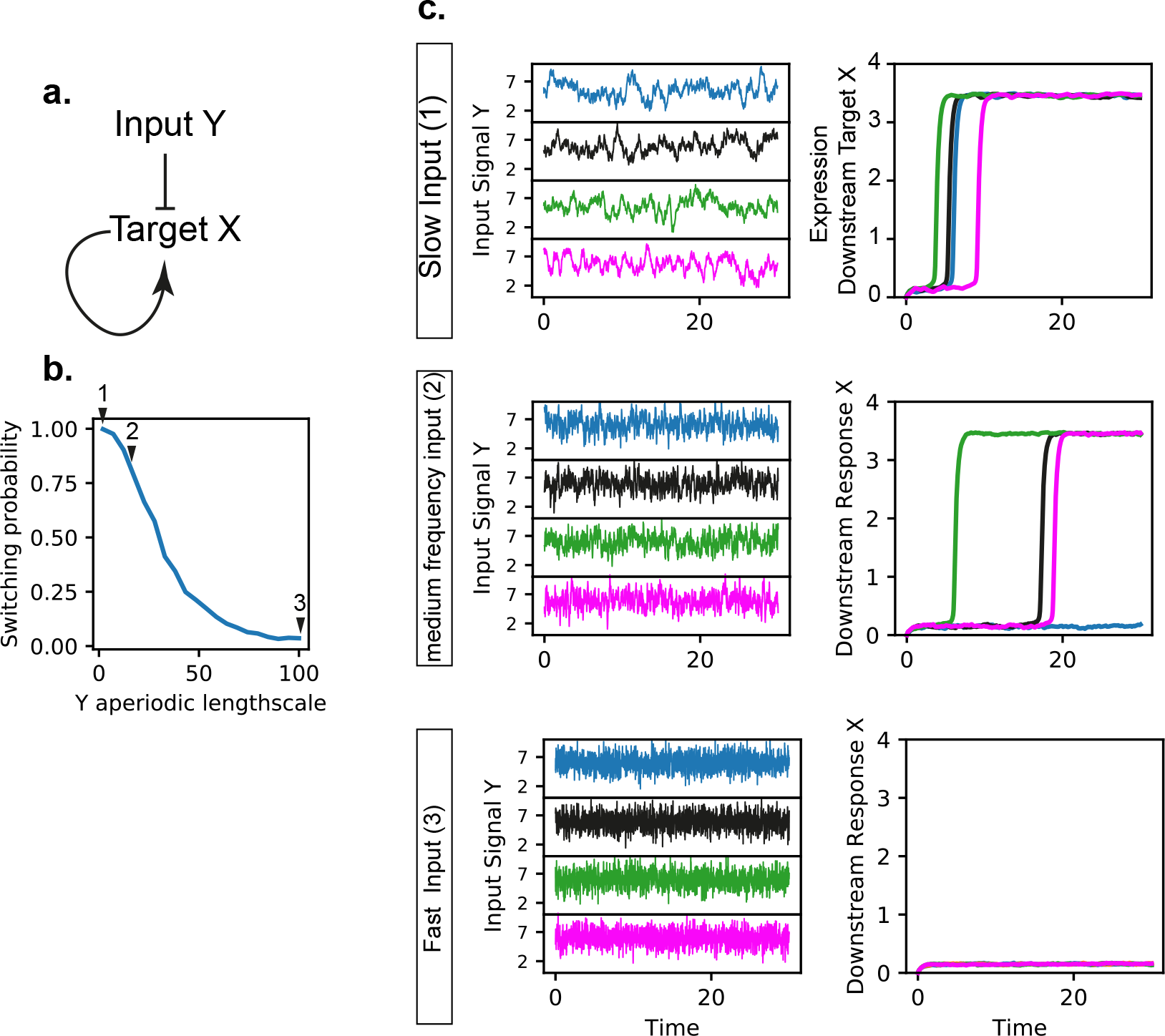
Changes in fluctuation dynamics can affect downstream target expression. **(a)** Network motif representing the interaction between a repressing gene Y acting as input onto a downstream target gene X; this motif represents a minimal network interaction that is sensitive to the aperiodic lengthscale. **(b)** Probability that the downstream target X switches to high expression from an initial off state; mathematical modelling shows that the probability decreases from one to zero as the aperiodic lengthscale in the dynamics of the repressing gene Y increases. **(c)** Example of gene expression dynamics of Y and X for different scenarios corresponding to slow, medium-frequency, and fast input, as quantified by aperiodic lengthscale levels highlighted in **(b**-arrows); multiple stochastic examples are shown for each scenario (left panel), and matching dynamics of X and Y are presented (right panel) in corresponding colours between the two panels; for increasing aperiodic lengthscales of Y, time to activation of X increases until X does not turn on within the depicted observation window.

Assuming that the gene X is initially not expressed, we found that it can switch towards high expression, depending on the dynamics of the upstream factor Y (Figure 5b). The probability to switch towards the high X expression within a finite observation window decreased with increasing aperiodic lengthscale of Y. For slowly varying input Y (Figure 5b,c; case 1), the probability that the gene X is turned on within the time window is one. For faster varying input (Figure 5b,c; case 2), the waiting time before the gene switches to high expression is increased, and in individual cases the switch may not happen within the observation time window. For a quickly varying input (Figure 5b,c; case3), the probability that X expression turns on within the observation window is zero (see Online Methods-Mathematical modeling for details of model and parameters).

Thus, our mathematical model predicts that the loss of oscillatory expression and the increased noise in the form of fast fluctuations in her6 expression can impede the upregulation of downstream genes that mediate a cell state transition, such as pro-neural genes.

### Her6 expression transcends cell state transitions; changes in Her6 dynamics affect cell state progression

To test this mathematical prediction we sought to determine the effect of altered Her6 dynamics on downstream targets that would normally be expressed when Her6 is downregulated. Late pro-neural transcription factors such as basic helix-loop-helix gene NeuroD4 is known to be downstream and regulated by Her/Hes family members ^32,33^; therefore we looked at the expression of NeuroD4, known as a marker of neuronal commitment. As predicted from the mathematical model, we frequently observed a decrease in the expression of *neuroD4* in MBSm embryos (Fig. 6a and b).

**Figure 6.**
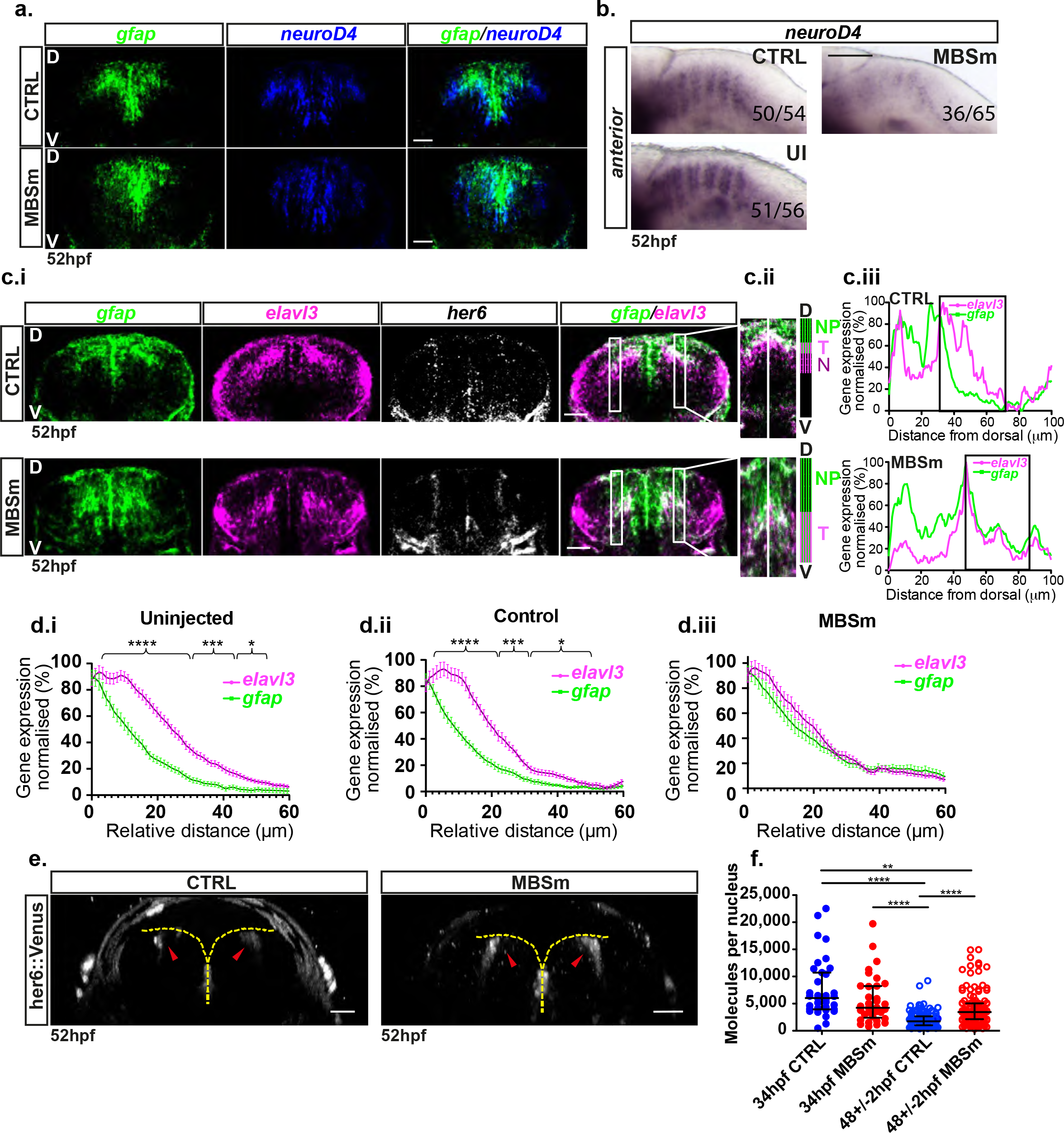
Effect of Her6 on downstream targets and cell fate decisions in the absence of miR-9 regulation. **(a)** Transverse sections of double F-WMISH for *gfap* (green) and *neuroD4* (blue) in the hindbrain (r6) of control (top) and MBSm (bottom) embryos at stage 52hpf; D: dorsal; V: ventral; scale bar 30μm. **(b)** Chromogenic WMISH for *neuroD4* in uninjected (UI); 51/56 embryos (91%) show normal expression intensity, control (CTRL); 50/54 embryos (93%) show normal expression intensity and mutant (MBSm); 36/65 embryos (55 %) show normal expression intensity, 45% were showing weak or/and abnormal expression; longitudinal view, scale 100μm. **(c.i)** Triple F-WMISH for *gfap* (green) and *elavl3* (magenta) and *her6* (grey) in the hindbrain (r6) of CTRL (top) and MBSm (bottom) embryos at 52hpf; scale bar 30μm. **(c.ii)** Magnification of inset from merged *gfap/elavl3* in **(c.i)** showing CTRL (top) and MBSm (bottom); annotations: neural progenitor zone, NP=*gfap(+)/elavl3(−)*; transition zone, T=*gfap(+)/elavl3(+)* and neurogenic zone, N=*gfap*(−)/*elavl3(+)*. **(c.iii)** Normalized intensity mean of *elavl3* and *gfap* along the DV axis spanning the: (CTRL: NP, T and N zones) and (MBSm: NP and T zones); region of interest (ROI) delineates high *elavl3* vs *gfap in* CTRL (T and N zones) while in MBSm only the T zone is observed. **(d)** Normalized mean of *elavl3* and *gfap* intensities in ROI observed in uninjected (5 embryos, 62 slices), control (5 embryos, 35 slices) and MBSm (8 embryos, 62 slices) conditions; bars represent mean and SEM; multiple t-test with Benjamini, Krieger and Yekuteli discovery. **(e)** Her6::Venus expression (red arrows) in hindbrain (r6) of live CTRL and MBSm embryos at 52hpf; edge of ventricular zone shown in yellow; transversal view; scale 30μm. **(f)** Her6::Venus protein abundance in control versus MBSm homozygous embryos at 34hpf vs 48hpf; bars indicate median and interquartile range of (34hpf CTRL: one embryo, 36 cells), (34hpf MBSm: one embryo, 35 cells), (48±2hpf CTRL: 6 embryos, 147 cells) and (48±2hpf MBSm: 5 embryos, 124 cells); statistical tests indicate Kruskal-Wallis with Dunn’s multiple comparison. Significance: p<0.05*, p<0.01**, p<0.001***, p<0.0001****.

To gain insight into a resultant phenotype, first we characterised in more detail the expression of *her6* in relation to cell states in the hindbrain. Triple F-WMISH staining was performed in embryos at 34hpf (suppl. Fig. S5) and 52hpf (Fig. 6c.i, top panel) including a progenitor marker, *gfap* and a neuronal differentiation marker, *elavl3*. *Elavl3* is expressed in neural cells early in their path to differentiation and is switched off in more basally located mature neurons ^25^. As expected, e*lavl3* and *gfap* are expressed in largely non-overlapping regions found in the dorsal-ventral axis of the hindbrain (Fig. 6c.i-top panel). However, a band of cells that co-express *gfap* and *elavl3* were identified, which we propose reflect a transitory state (Fig. 6.c.ii-top panel, transition zone T), as *gfap* expressing progenitors progress to *elav3* expressing early differentiating neurons (Fig. 6.c.ii-top panel, neurogenic zone N). The column of *her6* expression spans these domains, suggesting that it is expressed in progenitors (*gfap*(+)/*elavl*(−)), early differentiating neurons (*gfap*(−)/*elav3*(+)) as well as in cells of the identified transitory state (*gfap*(+)/*elavl3*(+)). *Her6* was not detected in cells located more basally, suggesting that it is downregulated along the neuronal differentiation pathway and switched off in mature neurons (Fig. 6c.i, top panel).

To determine if the changes in dynamic expression of Her6 have phenotypic consequences, F0 MBSm embryos were generated (Online Methods-microinjection and genotyping) and investigated by triple F-WMISH. In transverse sections, this showed a dorso-ventral expansion in the *gfap*(+) expressing progenitors and the *gfap*(+)/*elavl3*(+) transitory progenitors in MBSm embryos compared to control at 52hpf (Fig. 6c.i-ii, bottom vs top panel). This was accompanied by a decrease in the *gfap*(−)/e*lavl3*(+), suggesting that early differentiating neurons were not present in MBSm (Fig. 6c.ii, bottom panel). Quantitative analysis of *elavl3* and *gfap* levels in the Her6 expressing region, encompassed T and N zones in control and T zone only in MBSm, (Fig. 6c.ii-iii and 6d.i-iii), confirming the observed phenotype (Fig. 6c.i and Suppl Fig. S6b). We did not observe a phenotype at 28hpf when comparing CTRL to MBSm embryos (Suppl Fig. S6a.i-ii); this is consistent with the late miR-9 expression and its modulation of neurogenesis in late hindbrain development.

We also analysed Her6::Venus protein levels in live embryos by confocal fluorescence imaging (Fig. 6e, red arrowhead) and FCS quantitation of absolute nuclear abundance (Fig. 6f). Both methods showed that the decline in Her6 protein that normally takes place during development does not take place to the same degree in MBSm embryos, indicating that cells have not made the transition to differentiation. This is consistent with single cell Her6::Venus intensity traces observed using time lapse between 34hpf and 46hpf (Suppl. Fig. S6d.i-ii).

Taken together, these findings suggest that in the absence of miR-9 regulation, oscillatory expression of Her6 is reduced and noise is concomitantly increased, resulting in cells failing to downregulate Her6 and to progress through their normal cell state transition from progenitor to neuron

## Discussion

We have used the power of Zebrafish as an experimental system that combines live imaging with experimental perturbation, to interrogate the molecular dynamics of cell state transitions during neural development and their functional significance. We have focused on Her6, a key transcriptional repressor that belongs to a family of Her/Hes genes that have been shown to oscillate during vertebrate somitogenesis and mammalian neural development, and are essential for these processes ^13–15 7^

Using an endogenous CRISPR mediated knock-in fluorescent tag, we report for the first time the dynamics of a Her/Hes family member as they occur in real time, at the single cell level and in intact neural tissue. Our work was carried out in homozygous reporter Zebrafish, thus, the reported dynamics reflect what the cells experience. We found that Her6 expression undergoes a transition from irregular fluctuations (“noisy” expression) to oscillations with a dominant ultradian periodicity of approximately 2h as neurogenesis proceeds. This is consistent with our recent “ex vivo” tissue findings in the mouse spinal cord, where noise primes oscillatory expression by allowing the system to cross a bifurcation boundary which separates noisy from oscillatory expression ^12^. We have argued that this is an example of a beneficial use of noise in a biological system ^12^ joining other examples for the benefits of noise ^34–38 39^.

In Zebrafish, we were able to show that the transition switch from noisy to oscillatory dynamics coincides temporally with the onset of miR-9 expression, which has been previously proposed to be a “tuner” of Hes1 dynamics in the cultured mouse cells ^18 20 19,27^. Using CRISPR mediated knockdown of the miR-9 binding site in the her6 3’UTR, we functionally tested the role of miR-9 in modulating Her6 dynamics in vivo. An important finding of the present work is that miR-9 is needed for the oscillatory behavior of Her6 to emerge. Without the influence of miR-9, Her6 expression does not evolve away from the “noisy” into the oscillatory regime during development and moreover, in the absence of miR-9 there is an increase in high frequency noise in Her6 dynamics. We find that in the presence of such increased noise, neural progenitor cells are unable to make a transition to differentiation. This is accompanied by a failure of the natural reduction of Her6 protein levels during development and a concominant failure to upregulate early neuronal genes which are reported to be Her/Hes targets. The phenotype is consistent with previous static reports on the effect of miR-9 knock-down in Zebrafish ^17^ and Xenopus ^30 18^ but the dynamic nature of our findings further show the importance of oscillations in allowing the cells to make cell state transitions.

Moreover, our findings suggest an important role of noise optimisation in developmental transitions. The expression noise that we report here is characterised by an increase in frequency and as there was no pre-existing framework on the impact of such noise on gene expression, we developed a new computational model that allowed us to test its impact on downstream targets. We found that high frequency noise on Her6 expression prevents the activation of downstream pro-neural targets. This fits well with experimental observations whereby the proneural neuroD4 expression is decreased in miR-9 binding mutants. Thus, while a certain amount of noise may be beneficial in allowing the transition to oscillatory expression to occur, too much noise impedes the system of making this progression. We conclude that noise can be either beneficial or detrimental and noise optimization mediated by miR-9 is functionally necessary for cell state transitions.

What is the origin of the increased Her6 protein expression noise in the absence of miR-9? While the answer to this question is not presently known, we can offer two explanations that are supported by the literature. The first is that in the absence of miR-9 the translation of Her6 is increased allowing more of the stochasticity in gene expression through to the protein level. This fits well on the known effects of microRNAs in reducing translation ^40–42 43^ and the increase in noise when translation is increased ^29 44^. The second is that the absence of miR-9 decreases the robustness of the miR-9/Her6 network. This would happen if both miR-9 and Her6 (which normally repress each other, ^17 30^) respond to the same upstream signaling, forming an incoherent feedfoward loop. In such networks the negative influence of miR-9 to Her6 could be utilized to even out fluctuations in that signaling, as exemplified by other similar cases (reviewed in ^45^). Indeed, in support of this scenario it has been shown that both miR-9 and Her6 respond to upstream Notch signaling ^17^. Both of these scenarios converge in the negative regulation of her6 by miR-9 and further enrich the proposed roles of microRNAs in generating thresholds or dampening oscillations in other systems ^46 47^.

In conclusion, we have shown here that Her6 expression is dynamic in the tissue environment and that Her6 oscillations are facilitated by miR-9 mediated noise optimization; such oscillations are observed in progenitor cells but are functionally important for cells to transition to differentiation.

## Methods

### Research Animals

Animal experiments were performed under UK Home Office project licenses (PFDA14F2D) within the conditions of the Animal (Scientific Procedures) Act 1986. Animals were only handled by personal license holders.

### Generation of Her6::Venus knock in line

We used CRISPR/Cas9 technology combined with a DNA donor containing the Venus fluorescent protein to generate the *Her6::Venus* knock-in line (Fig. 1c).

We first identified the single guide RNA (sgRNA) to target *her6* exon 4, at the stop codon area using Addgene webpage, http://www.addgene.org/crispr/reference/#protocols, two or more software were utilised to choose the top scored sgRNAs, based in high efficiency and low off-target effect (see Online Methods-Preparation of Cas9 and sgRNAs section). We experimentally tested the sgRNA by High Resolution Melt (HRM) (See Method-Microinjection and genotyping) and chose the sgRNA with highest efficiency.

Further, we designed a DNA donor with either arms as big as 1kb ^48^ the left arm (LA) contained *Her6* exon2_intron2_exon3_intron3_exon4 and the right arm (RA) *Her6* exon4, we destroyed the sgRNA target site by inserting *linker_Venus* within. In order to generate the reporter controlled by endogenous Her6 expression we deleted the STOP codon from *Her6 gene* and the ATG codon from *Venus*. To avoid the inherent toxicity of linear DNA injection in fish, Two CRISPR target sites flanked the DNA donor thus; the fragment can be injected as a circular plasmid in pCRII vector but can be linearised in the cells to provide the linear template for the DNA repair once the Cas9-nls protein is translated from the co-injected mRNA ^49^ (See Online Methods-Molecular cloning).

Next, we co-injected the DNA donor with the sgRNA and Cas9nls mRNA. Later, we identified the Her6::Venus F0 adult fish carrying germline transmission following the method described online (preparation of Cas9nls and sgRNA in Online Methods-Microinjection and genotyping).

### Statistical testing

Comparative analysis between embryos at multiple conditions were carried out in Graphpad Prism 8.0. Quantitative data is presented per condition as box plot analysis with bars indicating median and interquartile range (5% and 95%) and statistically significant conditions reported for p values ≤ 0.05. Comparisons between normal distributed conditions were performed using one-way ANOVA with Bonferroni multiple comparisons test. Protein abundance from fluorescence correlation spectroscopy was analysed using Kruskal-Wallis with Dunn’s multiple comparison correction test. Dynamic parameters at multiple stages were presented as violin plots (bars: median and interquartile range; violin shape: distribution) and compared using a Mann-Whitney two-tailed test. Dynamic parameters were compared between control and mutant embryos from paired median values per experiment using a paired Student t-test with two-tail significance. Linear correlations with z position were tested using Pearson’s correlation coefficient. Non-linear correlations between signals collected from the same cells were tested using Spearman’s rank correlation coefficient. Intensity levels observed across the DV axis were adjusted to the same scale by linear interpolation and compared between conditions using multiple t-tests with Benjamini, Krieger and Yekuteli discovery, Q=1.

### Code and data availability

Detection of oscillators, aperiodic lengthscale and frequency analysis (Online Methods-Periodicity Analysis and Frequency Analysis) were performed using custom MATLAB 2018a routines. Mathematical model of network motif (Online methods-Mathematical Model) was implemented in Python. All code will be deposited in GitHub and is available upon request. Single cell Her6::Venus, mKeima-H2B raw and processed intensity traces are available upon request from the corresponding authors.

## Acknowledgements

We are grateful to Prof. Jon Clarke for help establishing transversal imaging in live Zebrafish and Prof Magnus Rattray for his continued support with statistical analysis of timeseries.We kindly thank Dr. Guilherme Costa, Dr Cerys Manning and Dr Anzy Miller for advice and discussions. The authors would also like to thank the Biological Services Facility and the Bioimaging Facilities of the University of Manchester for technical support. This work was supported by a Wellcome Trust Senior Research Fellowship to NP (090868/Z/09/Z). The funders had no role in study design, data collection and analysis, decision to publish, or preparation of the manuscript.

## Bibliography

1. Sagner, A. et al. Olig2 and Hes regulatory dynamics during motor neuron differentiation revealed by single cell transcriptomics. PLoS Biol 16, e2003127 (2018).

2. Farrell, J.A. et al. Single-cell reconstruction of developmental trajectories during zebrafish embryogenesis. Science 360(2018).

3. Levine, J.H., Lin, Y. & Elowitz, M.B. Functional roles of pulsing in genetic circuits. Science 342, 1193–200 (2013).

4. Shimojo, H., Ohtsuka, T. & Kageyama, R. Oscillations in notch signaling regulate maintenance of neural progenitors. Neuron 58, 52–64 (2008).

5. Imayoshi, I., Shimojo, H., Sakamoto, M., Ohtsuka, T. & Kageyama, R. Genetic visualization of notch signaling in mammalian neurogenesis. Cell Mol Life Sci 70, 2045–57 (2013).

6. Shimojo, H. et al. Oscillatory control of Delta-like1 in cell interactions regulates dynamic gene expression and tissue morphogenesis. Genes Dev 30, 102–16 (2016).

7. Kageyama, R., Shimojo, H. & Ohtsuka, T. Dynamic control of neural stem cells by bHLH factors. Neurosci Res 138, 12–18 (2019).

8. Sonnen, K.F. et al. Modulation of Phase Shift between Wnt and Notch Signaling Oscillations Controls Mesoderm Segmentation. Cell 172, 1079–1090 e12 (2018).

9. Nandagopal, N. et al. Dynamic Ligand Discrimination in the Notch Signaling Pathway. Cell 172, 869–880 e19 (2018).

10. Hatakeyama, J. et al. Hes genes regulate size, shape and histogenesis of the nervous system by control of the timing of neural stem cell differentiation. Development 131, 5539–50 (2004).

11. Novak, B. & Tyson, J.J. Design principles of biochemical oscillators. Nat Rev Mol Cell Biol 9, 981–91 (2008).

12. Manning, C.S., Biga, V., Boyd, J., Kursawe, J., Ymisson, B., Spiller, D. G., Sanderson, C. M., Galla, T., Rattray M., Papalopulu N. Quantitative, real-time, single cell analysis in tissue reveals expression dynamics of neurogenesis. bioRxiv (2018).

13. Soroldoni, D. & Oates, A.C. Live transgenic reporters of the vertebrate embryo’s Segmentation Clock. Curr Opin Genet Dev 21, 600–5 (2011).

14. Delaune, E.A., Francois, P., Shih, N.P. & Amacher, S.L. Single-cell-resolution imaging of the impact of Notch signaling and mitosis on segmentation clock dynamics. Dev Cell 23, 995–1005 (2012).

15. Webb, A.B. et al. Persistence, period and precision of autonomous cellular oscillators from the zebrafish segmentation clock. Elife 5(2016).

16. Leucht, C. et al. MicroRNA-9 directs late organizer activity of the midbrain-hindbrain boundary. Nat Neurosci 11, 641–8 (2008).

17. Coolen, M., Thieffry, D., Drivenes, O., Becker, T.S. & Bally-Cuif, L. miR-9 controls the timing of neurogenesis through the direct inhibition of antagonistic factors. Dev Cell 22, 1052–64 (2012).

18. Bonev, B., Stanley, P. & Papalopulu, N. MicroRNA-9 Modulates Hes1 ultradian oscillations by forming a double-negative feedback loop. Cell Rep 2, 10–8 (2012).

19. Phillips, N.E. et al. Stochasticity in the miR-9/Hes1 oscillatory network can account for clonal heterogeneity in the timing of differentiation. Elife 5(2016).

20. Goodfellow, M., Phillips, N.E., Manning, C., Galla, T. & Papalopulu, N. microRNA input into a neural ultradian oscillator controls emergence and timing of alternative cell states. Nat Commun 5, 3399 (2014).

21. Kesavan, G., Chekuru, A., Machate, A. & Brand, M. CRISPR/Cas9-Mediated Zebrafish Knock-in as a Novel Strategy to Study Midbrain-Hindbrain Boundary Development. Front Neuroanat 11, 52 (2017).

22. Zhou, M. et al. Comparative and evolutionary analysis of the HES/HEY gene family reveal exon/intron loss and teleost specific duplication events. PLoS One 7, e40649 (2012).

23. Schwanhausser, B. et al. Global quantification of mammalian gene expression control. Nature 473, 337–42 (2011).

24. Pasini, A., Henrique, D. & Wilkinson, D.G. The zebrafish Hairy/Enhancer-of-split-related gene her6 is segmentally expressed during the early development of hindbrain and somites. Mech Dev 100, 317–21 (2001).

25. Lyons, D.A., Guy, A.T. & Clarke, J.D. Monitoring neural progenitor fate through multiple rounds of division in an intact vertebrate brain. Development 130, 3427–36 (2003).

26. Zannino, D.A., Downes, G.B. & Sagerstrom, C.G. prdm12b specifies the p1 progenitor domain and reveals a role for V1 interneurons in swim movements. Dev Biol 390, 247–60 (2014).

27. Tan, S.L., Ohtsuka, T., Gonzalez, A. & Kageyama, R. MicroRNA9 regulates neural stem cell differentiation by controlling Hes1 expression dynamics in the developing brain. Genes Cells 17, 952–61 (2012).

28. Phillips, N.E., Manning, C., Papalopulu, N. & Rattray, M. Identifying stochastic oscillations in single-cell live imaging time series using Gaussian processes. PLoS Comput Biol 13, e1005479 (2017).

29. Kaern, M., Elston, T.C., Blake, W.J. & Collins, J.J. Stochasticity in gene expression: from theories to phenotypes. Nat Rev Genet 6, 451–64 (2005).

30. Bonev, B., Pisco, A. & Papalopulu, N. MicroRNA-9 reveals regional diversity of neural progenitors along the anterior-posterior axis. Dev Cell 20, 19–32 (2011).

31. d’Eysmond, T., De Simone, A. & Naef, F. Analysis of precision in chemical oscillators: implications for circadian clocks. Phys Biol 10, 056005 (2013).

32. Park, S.H. et al. Zath3, a neural basic helix-loop-helix gene, regulates early neurogenesis in the zebrafish. Biochem Biophys Res Commun 308, 184–90 (2003).

33. Bae, Y.K., Shimizu, T. & Hibi, M. Patterning of proneuronal and inter-proneuronal domains by hairy- and enhancer of split-related genes in zebrafish neuroectoderm. Development 132, 1375–85 (2005).

34. Balazsi, G., van Oudenaarden, A. & Collins, J.J. Cellular decision making and biological noise: from microbes to mammals. Cell 144, 910–25 (2011).

35. McDonnell, M.D. & Ward, L.M. The benefits of noise in neural systems: bridging theory and experiment. Nat Rev Neurosci 12, 415–26 (2011).

36. Hanggi, P. Stochastic resonance in biology. How noise can enhance detection of weak signals and help improve biological information processing. Chemphyschem 3, 285–90 (2002).

37. McDonnell, M.D. & Abbott, D. What is stochastic resonance? Definitions, misconceptions, debates, and its relevance to biology. PLoS Comput Biol 5, e1000348 (2009).

38. Moss, F., Ward, L.M. & Sannita, W.G. Stochastic resonance and sensory information processing: a tutorial and review of application. Clin Neurophysiol 115, 267–81 (2004).

39. Paulsson, J., Berg, O.G. & Ehrenberg, M. Stochastic focusing: fluctuation-enhanced sensitivity of intracellular regulation. Proc Natl Acad Sci U S A 97, 7148–53 (2000).

40. Filipowicz, W., Bhattacharyya, S.N. & Sonenberg, N. Mechanisms of post-transcriptional regulation by microRNAs: are the answers in sight? Nat Rev Genet 9, 102–14 (2008).

41. Beilharz, T.H. et al. microRNA-mediated messenger RNA deadenylation contributes to translational repression in mammalian cells. PLoS One 4, e6783 (2009).

42. Bazzini, A.A., Lee, M.T. & Giraldez, A.J. Ribosome profiling shows that miR-430 reduces translation before causing mRNA decay in zebrafish. Science 336, 233–7 (2012).

43. Zinovyev, A., Morozova, N., Gorban, A.N. & Harel-Belan, A. Mathematical modeling of microRNA-mediated mechanisms of translation repression. Adv Exp Med Biol 774, 189–224 (2013).

44. Raj, A. & van Oudenaarden, A. Nature, nurture, or chance: stochastic gene expression and its consequences. Cell 135, 216–26 (2008).

45. Ebert, M.S. & Sharp, P.A. Roles for microRNAs in conferring robustness to biological processes. Cell 149, 515–24 (2012).

46. Mukherji, S. et al. MicroRNAs can generate thresholds in target gene expression. Nat Genet 43, 854–9 (2011).

47. Kim, D., Grun, D. & van Oudenaarden, A. Dampening of expression oscillations by synchronous regulation of a microRNA and its target. Nat Genet 45, 1337–44 (2013).

48. Zu, Y. et al. TALEN-mediated precise genome modification by homologous recombination in zebrafish. Nat Methods 10, 329–31 (2013).

49. Irion, U., Krauss, J. & Nusslein-Volhard, C. Precise and efficient genome editing in zebrafish using the CRISPR/Cas9 system. Development 141, 4827–30 (2014).

