## Supplemental Methods, supplemental table and supplemental figures for "miR-9 mediated noise optimization of the her6 oscillator is needed for cell state progression in the Zebrafish hindbrain"

#### Online Methods

##### Molecular cloning

The DNA Donor was constructed in the pCRII vector (Life Technologies). The Her6-LA, Her6-RA and linker-Venus were generated by PCR, using genomic DNA of wild-type (AB) fish and her1:her1-linker-Venus (Delaune et al., 2012) respectively, using specific primers (see suppl. Table1). Primers 1 and 4 were containing the sgRNA target site and primer 5 wasn't including the start coding sequence for Venus. The PCR products were cloned into pCRII vector followed by sequential subcloning steps to assemble the DNA donor, first we used SalI/SpeI to subclone LA into pCRII_linker-Venus then it was used KpnI to subclone linker-Venus_LA into pCRII _RA.

The pCS2+ mKeima-H2B was generated by sequential PCR, restriction enzyme treatment and subcloning steps, using the plasmid mKeima-Red-N1 (Addgene #54597) as template. The pCS2+ HA::Her6 and pCS2+ Her6::Venus were generated by sequential PCR and subcloning steps, using as a template cDNA obtained by RT from embryonic mRNA (see suppl. Table1 for respective primer set).

##### Preparation of Cas9nls and sgRNAs

The Cas9nls mRNA we generate from pT3TS-nls-zCas9-nls plasmid obtained from Addgene #46757 following the protocol described by Li-En Jao [1]. The Cas9nls protein was obtained from New England Biolabs M0641M.

The sgRNA target sites were identified using the CRISPRdirect (http://crispr.dbcls.jp/) and Target Finder (Feng Zhang lab. http://crispr.mit.edu/).

For Her6::venus knock-in the selected oligonucleotides (suppl. Table 1) were annealed and cloned into pT7-gRNA plasmid #46759 (Addgene), following the protocol described by Li-En Jao [1]. The correct clones were linearised with BamHI and transcription of sgRNA was carried out using MEGAshortscript T7 kit (Ambion/Invitrogen) with 100-400 ng of purified linearised DNA following

the manufacturer's instructions. The sgRNA was purified using MEGAclear™ Transcription Clean-Up Kit.
For Her6 miR-9 binding site (MBS) mutation, sgRNAs were generated following CRISPRscan protocol [2]) using the oligonucleotides (25 or 26) described in suppl. Table1. The PCR fragments were purified using Qiagen columns and transcriptions were carried out as described above.

**Microinjection and genotyping**

To generate the Her6::Venus knock-in (Ki), one-cell stage wild type AB Zebrafish embryos were injected with ~1 nl of a solution containing 150 ng/µl Cas9nls mRNA, 200 ng/µl sgRNA, 20 ng/µl circular DNA donor in 0.05% phenol red.
To generate MBS mutation, one-cell stage Her6::Venus Ki embryos were injected with ~1 nl of a solution containing 185 ng/µl Cas9nls protein, 125 ng/µl sgRNA, 40 ng/µl caax-mRFP mRNA, 40 ng/µl mKeima-H2B mRNA in 0.05% phenol red.
To evaluate efficiency of sgRNA, genomic DNA was extracted from 3-4dpf embryos using 50 µl NP lysis buffer per embryo (10mM Tris pH8, 1 mM EDTA, 80mM KCl, 0.3 % NP40 and 0.3% Tween) and 0.5 µg/µl Proteinase K (Roche) for 3-4 hrs at 55°C, 15 min at 95°C and then stored at 4°C. Then, High Resolution Melt (HRM) was performed using Melt Doc kit following manufacturer instructions, specific primers set were used according Her6::Venus Ki or MBS mutation (See suppl. Table1).
To assess homology direct repair (HDR) in F0 or F1 progeny, genomic extraction was performed as described above followed by PCR and agarose gel (See suppl. Table1). The PCR products were cloned into pCRII vector and sequenced. It is relevant to mention that the repair of the Her6-RA wasn't perfect as it included part of the vector that was used to generate the DNA donor; nevertheless we expected this insertion to not affect the protein expression, and dynamics, as the Her6-RA included the 3'UTR and Her6 polyadenylation site (suppl. Fig. S1C.i.).

To identify F1 progeny with germ line transmission (GLT) 3-5dpf embryos were fin clipped following the protocol described by Robert Wilkinson (Wilkinson R. et al., 2013) with modifications. Sylgard (Sigma, Cat # 761028)-coated 10 cm dish was prepared for dissections. Embryos were placed into Sylgard-coated dish containing E3 medium with 0.1% Tricaine (Sigma, UK) and 2% BSA (Sigma, UK). Once clipped the fin the embryo was transferred to E3 medium and the biopsy was transferred to PCR tube for genomic extraction. Genomic extraction was carried out in 10 µl volume using Phire Animal Tissue Direct PCR kit (Thermo scientific, Cat # F-140WH). 2µl of the supernatant was used for 10 µl qPCR using primers N9 and N12. Derivative of the Melting Curve was used to identify the positive GLT fish.

**Whole mount chromogenic and fluorescence *in situ* hybrisation and sectioning**

Chromogenic *in situ* hybridisation was carried out as described by Christine Thisse [3]. Multicolour fluorescence *in situ* hybridisation was developed using tyramide amplification after addition of probes and antibodies conjugated to horseradish peroxidase [4]. RNA probes for Elavl3, GFAP, Neurod4, Her6, Venus, pri-miR-9-4 were PCR amplified and cloned into pCRII vector using the primers in suppl. Table 1. Eplin probe was generated from a plasmid kindly gifted by Andrew Oates. miR-9 LNA 3'5'Dig probe was purchased from Exiqon.

Sections were obtained as described in Dubaissi [5] with modifications. Embryos were embedded in 25% fish gelatine for a minimum of 24 hrs. 18 μm thickness section were collected and transferred onto superfrost glass slides. The slides were air dried for 6 hrs under fume hood and washed for 2 min in PBS only before mounting.

**Protein Half-life**

One-cell stage wild-type embryos were injected with 80pg of Her6::Venus or HA::Her6 mRNA. Protein half–life was performed using 200μM Cycloheximide and incubation started at 128-256 cells, development stage. Pools of 20 embryos were collected at 0, 5, 10, 15 and 20 min. Samples were lysated with 2 μl/emb of Ginzburg Fish Ringer lysis buffer (110mM NaCl, 3.35mM KCl, 25mM CaCl2 and 2.4mM NaHCO3) and washed twice with 500 μl of Ginzburg Fish Ringer wash buffer (110mM NaCl, 3.5mM KCl, 2.7mM CaCl2 and 10mM Tris pH 8.8). The pellet was resuspended in 2μl/emb of 1x Laemmli buffer. Western blots were performed using 4-20% Tris-glycine acrylamide gels (NuSep), Trans-Blot Turbo Midi Nitrocellulose Transfer Pack (Bio-rad) and developed with Pierce ECL substrate (ThermoFisher Scientific). Antibodies used were anti-GFP (mouse Roche 1181446001), anti-HA-HRP (rat monoclonal Roche 2013819) and anti-alpha-tubulin (clone DM1A Sigma T9026).

**Fluorescent correlation spectroscopy**

For FCS experiments embryos were mounted in a customized metallic device, microscope slide shape hollowed in the middle in order to fix cover slip in either side. The embryos were mounted with 1% Low-melting agarose (Sigma) with the region of interest close to the cover slip.

Snapshot images were collected with Zeiss LSM880 microscope with a C-Apochromat 40x 1.2 NA water objective. FCS signals were collected inside single nuclei in dorsal region of the hindbrain in intact embryo. Venus (EYFP) fluorescence was excited with 514 nm laser light and emission collected between 517 and 570nm. Data from individual cell nuclei was collected using 5 x 5 s runs at 0.15 to 0.3% laser power which gave <10% bleaching and a suitable count rate ~1 kHZ counts per molecule (CPM). To obtain molecule number, autocorrelation curves were fit to a two-component diffusion model with triplet state using an optimization Toolbox based on the Levenberg-Marquardt algorithm with initial conditions assuming a ‘fast’ diffusion component 10x faster than the ‘slow’ component as described in Smyllie et al. [6].

Measurements collected from cells exhibiting large spikes/drops in count rate or with low CPM (<0.5 kHz), high triplet state (>50%), or high bleaching

(>10%) were excluded from the final results. Number and brightness analysis of the count rate showed a high correlation with molecule number obtained from autocorrelation curve fitting. The effective confocal volume (CV) had been previously determined with mean 0.57fL ± 0.11fL [7], allowing conversion from molecule number to concentration. Single-cell data of absolute protein number in the cell nucleus was obtained by adjusting concentration in CV to the average volumetric ratio between nuclear volume and confocal volume. Cell volumes were larger at 28hpf (379.32 ±26.39fL), compared to 34hpf (118.13 ±5.9fL). CTRL and MBSm at 34hpf showed no significant differences in volume (volumetric ratio CTRL vs MBSm=1.01).

**Live imaging**

For confocal microscopy live embryos were mounted in 1% low-melting agarose in glass-bottom dishes, which were subsequently filled with media supplemented with 0.0045% 1-phenyl-2-thiourea and 0.1% tricaine. Embryos were imaged using HCX IRAPO 25x0.95 water dipping objectives on a Leica TCS SP5 or TCS SP8 upright confocal microscopes, pinhole 2AU. Embryos were maintained at 28°C whilst imaging using an in-line solution heater and heated stage, both controlled by a dual-channel heater controller (Warner Instruments).

To study the overall Her6 dynamic during Hindbrain development the embryos were laterally mounted, images were collected every 10 min, 60 Z-stacks per time point and 1.3microns voxels depth. To study Her6 dynamic at single cell resolution the embryos stage 28, 30 or 34hpf were mounted with forebrain facing down (Fig. 3A) in order to collect a transversal view of the rhombomere 6, images were collected every 6 min, 12-15 Z-stacks per time point and 2.0 microns voxel depth, with x4 magnification.

**Cell tracking and Image pre-processing**

We used the arithmetic tool in Imaris to segment the Venus channel. First, the mKeima channel (nuclei signal) was substracted from the mRFP channel (membrane channel) to remove noise and autoflorescence signal coming from

apoptotic cells. This new channel was substracted to the Venus channel to better define the contour of cells expressing Her6::venus (suppl. Fig. S2a).
Long-term (above 3h) trends in the relative Her6/H2B signal are removed as described in Phillips et al. 2017 to generate the detrended Her6/H2B signal.
Single neural progenitor cells in rhombomere 6 were tracked in Imaris on the Her6::Venus substracted channel using the 'Spots' and 'Track over time' function. Tracking function used the Brownian motion algorithm. All tracks were manually curated to ensure accurate single-cell tracking.
To account for any photobleaching for Her6::Venus (suppl. Fig. S2b) we measured tissue background over time and calculated the linear decay per channel (Fig. S2b).
To account for any correlation between Her6::Venus and mKeima-H2B (suppl. FigS2c) we normalised Venus signal to mKeima (examples show in Fig. S3a,b) in order to correct for any global changes in transcription or translation in the cell or microscope anomalies.
Long-term (above 3h) trends in the relative Her6/H2B signal are removed as described in [8] to generate the detrended Her6/H2B signal.
We also investigated the relationship between fluorescence intensity and Z position in the tissue. As expected from imaging through tissue there was a small negative correlation of Her6::Venus and mKeima-H2B intensity associated with Z position, r = -0.114 and r = -0.148 respectively, when all cells and time-points were plotted (suppl. Fig. S2d-e). However the range of Z positions in a single cell 10-12 hour track was rarely greater than 25μm, therefore it is unlikely the fluctuations and oscillations in Her6::Venus are a result in changes in Z position.

**Mathematical model**

We implemented a mathematical model for the response of the expression of a target gene X to different dynamics of Her6 expression. The model allows calculating the response of the target gene to pre-specified traces of Her6 expression with different dynamic properties. The interactions governing the dynamics of the target gene form a minimal network that is sensitive to the timescale of Her6 fluctuations. We assumed that the downstream target X would be able to self-activate, which is consistent with the literature [9]. We expect other network motifs to have qualitatively similar properties. For example, the self-activation could appear as a consequence of repressing a repressing gene.In the following we provide a mathematical description of the model and explain different Her6 dynamics are generated.

The mathematical model for the response of the target gene X to Her6 fluctuations is implemented using

$$\frac{dX}{dt} = G_1\big(y(t)\big) + G_2(X) - \mu X \qquad (1)$$

where $X$ is the protein copy number of the target gene, $t$ is time, $y(t)$ is copy number of Her6 at time $t$ and $\mu$ is the protein degradation rate of the downstream target gene. The production functions $G_1(y)$ and $G_2(X)$ describe the repression of $X$ by Her6 and its auto-activation, respectively, by defining how the presence of Her6 or $X$ can influence $X$ production. We chose a production function $G_1(y)$ that returns a finite production rate, $k_1$, if Her6 copy numbers $y$ are much lower than a threshold of repression, and which leads to a production rate of 0 if Her6 copy numbers are much greater than this repression threshold. We denote the repression threshold by $y_0$, which allows us to define

$$G_1(y) = k_1 \frac{1}{1 + (\frac{y}{y_0})^n},$$

where $n$ is a Hill coefficient controlling steepness of the transition from $k_1$ to 0 as the Her6 concentration $y$ increases. The function $G_2(X)$ is defined in similar manner,

$$G_2(X) = k_2 \frac{1}{1 + (\frac{X}{X_0})^{-n}}.$$

This function leads to a production rate $k_2$ of $X$ if the number of $X$ molecules is much greater than the activation threshold $X_0$, and 0 if the number of $X$ molecules is much smaller than the activation threshold. Again, the steepness of the transition is regulated by the Hill coefficient $n$.

*Simulation of Her6 fluctuations*

We generated *in silico* Her6 dynamics by sampling traces from a OU process, which is introduced using the covariance function $K_{OU}$ in the main text. Varying the aperiodic lengthscale of the process can generate traces with different timescales of fluctuation, similar to the differences we see between the data of the MBS and the control experiments. In order to ensure non-negative values for these *in silico* traces, we added the Ornstein Uhlenbeck traces to an otherwise constant and non-negative signal. Once we had defined *in silico* Her6 dynamics $y(t)$ in this way, we used equation (1) to simulate the downstream response of X expression.

*Choice of Parameter Values*

Dynamical systems, such as the one illustrated in Figure 5 and described by equation (1), may exhibit different qualitative behaviors in different regions of parameter space. To generate 5, we manually selected parameters that place the model in a regime where it exhibits sensitivity to the timescale of the Her6 fluctuations. Specifically, *in silico* Her6 dynamics were generated by combining a constant expression at a level of 6 arbitrary units with a sample from an OU process for which we used a variance parameter of 1.73 and different values for $\alpha$ (case 1: $\alpha$=2, case 2: $\alpha$=15, case 3: $\alpha$ =100). For the dynamics of $X$, we used the parameters $k_1$ = 0.5, $k_2$ = 10, $y_0$=7.9, $X_0$=0.5, $n$ =4, $\mu$= 3. We implemented the model using a standard forward Euler scheme with a time step of 0.0015 and a simulation duration of 30 time units. Note, however, that the observed qualitative behavior of the model is not dependent on the specific parameter choices, and similar graphs to those in figure 5 can be generated with multiple different parameter combinations.

**Periodicity analysis**

The dynamic data analysis was performed using custom routines amended from the approach in [8] and code generated by Nick Phillips (deposited at https://github.com/ManchesterBioinference/GPosc) and using the GPML toolbox (details at http://gaussianprocess.org/gpml/code/matlab/doc/) [10], implemented in MATLAB R2015aSP1 . Specifically, we use Gaussian

Processes that can represent timeseries in terms of a *mean function*, capturing long term (above 3hrs) changes in the signal and *a covariance function* describing variations around the mean. Periodic and aperiodic dynamic activity is characterized by two competing covariance functions, one encoding *aperiodic and stochastic* fluctuations, $K_{OU}(\tau) = \sigma \exp(-\alpha_{OU}\tau)$ and another encoding *periodic and stochastic* activity through an additional wave term, $K_{OUosc}(\tau) = \sigma \exp(-\alpha_{OUosc}\tau) \cos(\beta\tau)$. The signal variance, $\sigma$ is included in both models and is related to amplitude. Since we discuss amplitude separately in relation to level (see LCOV), data is z-scored prior to analysis leading to $\sigma = 1$.

Stochasticity is described in both models by an exponential decay in correlation between subsequent peaks and parameterised by the rate of decay parameter$\alpha_{OU}$, aperiodic lengthscale and $\alpha_{OUosc}$ , periodic lengthscale for $K_{OU}$ and $K_{OUosc}$ respectively. In addition, the periodic covariance model includes a cos wave term with frequency $\beta$ and related to period by $P = 2\pi/\beta$. Parameters are inferred from data by maximum likelihood techniques where likelihood represents the probability of the observed data under the model and includes a technical noise term.

*Parameter estimation*

We have previously shown that fluorescence data acquired in 3D has a characteristic low signal-to-noise ratio (90%) which impacts log-likelihood, thus the estimation of multiple parameters required additional considerations for the analysis if periodicity {Manning, 2018 #151. Consistent with this, we define a flexible prior for the estimation of periodic lengthscale. This strategy constrains the values of lengthcale estimated under the OUosc model and increases the ability to discriminate between oscillatory and non-oscillatory cells. In the absence of sufficient samples to perform a robust joint log-likelihood approach, we revert to an experimental calibration of technical noise. Namely we determine the variance of technical noise to variance observed in an area of the tissue found outside the Her6(+) area but expressing mKeima-H2B (referred to as tissue background). This ensure that the covariance models do not fit spurious dynamics observed below the detection limit.

*False Discovery Rate*

We determine oscillators and non-oscillators using the false discovery rate approach in {Phillips, 2017 #153}. The statistic used is the log-likelihood ratio (LLR) which compares the likelihood of the periodic and aperiodic models such that a large LLR value indicates high probability of the timeseries being oscillatory while a value around 0 indicates high probability of aperiodic fluctuations. We used a strict 3% FDR and analysed the data per experiment.

**Frequency analysis**

Frequency analysis was performed using MATLAB R2015aSP1. The power spectrum reconstruction per conditions was generated using the routine *periodogram.m* evaluated at a fixed frequency range and averaged over the number of cells analysed per experiment. The single cell power spectral density (PSD) reconstruction was generated using *periodogram* with the option '*psd'* whereby the power is normalized by signal variance. We choose a cutoff between the ultradian region and the high frequency region corresponding to 40min. The % contribution of high frequency to the variance in the data is quantified by integrating the PSD across the high frequency region and dividing by the area under the curve.

**COV and LCOV**

Coefficient of variation (COV) denoting the ration between standard deviation and the sample mean was used to describe the variability of the signal. To account for the presence of long-term trends (above 3h), we analyse COV in a (local) sliding time window set to 1.4h (~ period) and refer to this measure as local COV or LCOV.

**Supplementary Table1.** Oligonucleotide sequences used for cloning, sgRNA, High Resolution Melt (HRM) and Genotyping.

| No. | NAME | Sequence 5’-3’ |
|---|---|---|
| 1 | her6 LA cloning forward | ggatccggcggccttggtagactccgagggaactatagtgcttcaagaagtgattag |
| 2 | her6 LA cloning reverse | agtcgaccctccactacctccccaaggccgccaaacggagtctgacg |
| 3 | Her6 RA cloning forward | tggtaccactccgaggagaaaaaaaaactcttaaaagac |
| 4 | her6 RA cloning reverse | ggcggccttggtagactccgaggccgcagtgtcagaaacaccgtatattaacatg |
| 5 | Linker_venus cloning forward | agggtcgactgctagcggtggagtgagcaagg |
| 6 | Linker_venus cloning reverse | aaggtaccttacttgtacagctcgtccatgccg |
| 7 | her6 CRISPR sgRNA forward | tagg**ggcggccttggtagactccg** |
| 8 | her6 CRISPR sgRNA reverse | aaac**cggagtctaccaaggccgcc** |
| 9 | her6 CRISPR HRM forward | gtctacgcaaacaattccaa |
| 10 | her6 CRISPR HRM reverse | cgctgaacaaagaaaacaagt |
| 11 | her6 CRISPR genotyping LA forward | atgcctgccgatatcatgga |
| 12 | her6 CRISPR genotyping LA reverse | tgaacagctcctcgcccttg |
| 13 | gfap probe cloning forward | attctcctccaccatggag |
| 14 | gfap probe cloning reverse | agtgaaggagatcttcttctg |
| 15 | elavl3 probe cloning forward | gtgcatcttcgtctacaacctg |
| 16 | elavl3 probe cloning reverse | acagataactgcatgtggtgg |
| 17 | neurod4 cloning probe forward | agtgaggcacgagacgcgctctg |
| 18 | neurod4 cloning probe reverse | ttattcgctgtaaatgctgctc |
| 19 | her6 probe cloning forward | tcgtcgacaagatgcctgccgatatcatgg |
| 20 | her6 probe cloning reverse | ccaaggccgccaaacggagtctgacg |
| 21 | venus probe cloning forward | agtggagggtcgactgctagcggtggc |
| 22 | venus probe cloning reverse | aaggtaccttacttgtacagctcgtccatgccg |
| 23 | Pri-miR-9-4 probe cloning forward | ttccacaagggtatcgatag |
| 24 | Pri-miR-9-4 probe cloning reverse | tattatatgagaaccacgtg |
| 25 | her6 CRISPRscan sgRNA MBS mutation | taatacgactcactatagggcgcattcaacatatcttgttttagagctagaa |
| 26 | her6 CRISPRscan sgRNA MBS control | taatacgactcactataggatctttggcatcacaacggttttagagctag aa |
| 27 | her6 MBS mutation HRM forward | tgctgtagtgtggaaccactag |
| 28 | her6 MBS mutation HRM reverse | tgtattgttgaattccgttcaatgc |
| 29 | mKeima BamHi cloning forward | cgggatccaccggtcgccaccatg |
| 30 | mKeima BspeI cloning reverse | aattccggaaccgagcaaagagtggcgtg |

**Supplementary Bibliography**


1. Jao, L.E., Wente, S.R. & Chen, W. Efficient multiplex biallelic zebrafish genome editing using a CRISPR nuclease system. *Proc Natl Acad Sci U S A* **110**, 13904-9 (2013).

2. Moreno-Mateos, M.A. *et al.* CRISPRscan: designing highly efficient sgRNAs for CRISPR-Cas9 targeting in vivo. *Nat Methods* **12**, 982-8 (2015).
3. Thisse, C. & Thisse, B. High-resolution in situ hybridization to whole-mount zebrafish embryos. *Nat Protoc* **3**, 59-69 (2008).
4. Lea, R., Bonev, B., Dubaissi, E., Vize, P.D. & Papalopulu, N. Multicolor fluorescent in situ mRNA hybridization (FISH) on whole mounts and sections. *Methods Mol Biol* **917**, 431-44 (2012).
5. Dubaissi, E., Panagiotaki, N., Papalopulu, N. & Vize, P.D. Antibody development and use in chromogenic and fluorescent immunostaining. *Methods Mol Biol* **917**, 411-29 (2012).
6. Smyllie, N.J. *et al.* Visualizing and Quantifying Intracellular Behavior and Abundance of the Core Circadian Clock Protein PERIOD2. *Curr Biol* **26**, 1880-6 (2016).
7. Bagnall, J. *et al.* Quantitative dynamic imaging of immune cell signalling using lentiviral gene transfer. *Integr Biol (Camb)* **7**, 713-25 (2015).
8. Phillips, N.E., Manning, C., Papalopulu, N. & Rattray, M. Identifying stochastic oscillations in single-cell live imaging time series using Gaussian processes. *PLoS Comput Biol* **13**, e1005479 (2017).
9. Helms, A.W., Abney, A.L., Ben-Arie, N., Zoghbi, H.Y. & Johnson, J.E. Autoregulation and multiple enhancers control Math1 expression in the developing nervous system. *Development* **127**, 1185-96 (2000).
10. Rasmussen, C.E. & Nickisch, H. Gaussian Processes for Machine Learning (GPML) Toolbox. *Journal of Machine Learning Research* **11**, 3011-3015 (2010).

#### Supplementary Figure Legends

**Figure S1. Generation and characterization of the Zebrafish knock-in reporter.**

**(a)** Diagram of experimental approach used to generate the Her6::Venus knock-in. **(b)** Amplicon generated by using primers 11 and 12, indicated by red arrowhead, obtained only when *venus* is inserted at C-terminus of *her6* gene. C-: uninjected embryos; C+: Her6::venus plasmid used as positive control. **(c.i)** Schematic of the Her6::Venus knock-in including a vector sequence inserted after the Her6 3'UTR. E: exon, **(c.ii)** Sequencing showing the correct DNA repair in exon1 of *her6* gene. **(c.iii)** Chromogenic WMISH for *venus* in Her6::Venus knock-in at 18 somites development (left); comparison of number of somites observed in homozygote, heterozygote and wild type embryos at 72hpf (right); bars represent mean and SD of (hom: 7 embryos), (het: 19 embryos) and (wt: 6 embryos). **(c.iv)** Pairwise comparison of protein abundance measured by FCS in the hindbrain of homozygous versus heterozygous embryos containing the Her6::Venus knock-in; data represents median per experiment from (hom: 4 embryos, 90 cells), (het: 4 embryos, 72 cells). (**d.i)** Genotyping by fin clipping at 3dpf, qPCR amplification **(d.ii),** followed by identification of positive fish by derivative of melt curve **(d.iii); (d.ii-iii:** ctrl(+) denotes positive PCR control; positive knock-in is marked by green arrow). **(d.iv)** Representative example of genotyping by fin clipping at 8wpf. **(e)** Half-life of HA-Her6 protein versus Her6::Venus protein measured in vitro by injection of mRNA in embryos; bars represent median and interquartile range from 3 biological repeats.

**Figure S2. Pre-processing of video data.**

**(a)** Raw images of hindbrain (r6) at 30hpf showing membrane marker: caax-mRFP (grey), nuclear marker: mKeima-H2B (magenta) and Her6::Venus (green); image processing steps included subtraction of nuclear from membrane marker to generate a caax-mRFP subtracted channel; this is then subtracted from Venus to produce a Her6::Venus segmented channel with enhanced separation between nuclei; scale bar 30µm. **(b)** Rate of bleaching observed as mean per experiment in the Venus and mKeima channels; error bars represent mean and SD per condition. **(c)** Spearman rank correlation coefficient calculated from Her6::Venus versus mKeima-H2B time series observed in the same cell; bars represent median and interquartile range of 35 cells from 3 embryos. **(d)** Visualisation of Her6::Venus intensity normalized per experiment versus z-position in all data collected; statistical p-value obtained form Pearson linear correlation test. **(e)** Visualisation of mKeima-

H2B intensity normalized per experiment verus z-position in all data collected; statistical p-value obtained from Pearson's linear correlation test.

**Figure S3. Single cell dynamics observed in Her6::Venus neural progenitor cells at different stages in development.** Representative examples of cells classified as oscillatory **(a)** and non-oscillatory **(b)** showing corresponding single cell time series of Venus::Her6 (panel 1), mKeima-H2B (panel 2), Her6 signal relative to H2B (panel 3) and detrended Her6/H2B signal (panel 4); dynamic statistics include: **(a-panel 4)** log-likelihood ratio (LLR), period and quality per cell and **(b-panel 4)** LLR (see Online Methods-Periodicity analysis).

**Figure S4. Mir-9 binding site (MBS) manipulation and frequency analysis.**
**(a)** Chromogenic WMISH of mir-9 using mir-9 LNA 5'-Dig observed at different stages during development; longitudinal view, anterior to the left.
**(b-left)** Schematic representation of experimental procedure used to mutate the Her6 mir-9 binding site; **(b-right)** high resolution melt graph obtained from: control (CTRL-blue) versus uninjected (UI-brown); and MBS mutation (MBSm-green) versus uninjected (UI-magenta). **(c)** Annealing of the MBS sequence obtained from F0 embryos injected with sgRNA to produce mutation and insertion (red); wild type sequence shown at top. **(d)** Power spectrum reconstruction from Her6/H2B detrended time series collected from CTRL versus MBSm embryos imaged starting from 34hpf; data represents population average of 14-15 cells per embryo per condition. **(e)** Heatmap representation of single cell power spectra normalized by variance (power spectral density, PSD) observed in single cells from CTRL versus MBSm embryos; single cell data corresponds to the population average included in **(d)**. **(f)** Quantification of the contribution of high frequency noise to single cell PSD data included in **(e)**; bars represent median and interquartile range of 14-15 cells per embryo per condition; statistical test represent Mann-Whitney with two-tail significance **($p<0.01$).

**Figure S5. Spatial localization of Her6 related to neural and progenitor markers. (top)** Representative examples of triple F-WMISH labelling of *gfap* (green), *elavl3* (magenta) and *her6* (grey) domain of expression. **(bottom)** Merged images indicating how the her6 expression domain overlaps with the progenitor zone (*gfap*(+)/*elavl3*(-)), transition zone (*gfap*(+)/*elavl3*(+)) and neurogenic zone (*gfap*(-)*/elav3l*(+)); embryo observed at 34hpf, transversal view; annotations denote dorsal (D) and ventral (V); scale bar 30µm; **(bottom-right panel)** schematic representation showing the *her6* domain spanning the progenitor, transition and neurogenic zones and with distance from dorsal.

**Figure S6. Effect of mir-9 binding site manipulation (MBSm) on expression of Her6 and downstream targets.**
**(a.i)** Triple F-WMISH for *gfap* (green), *elalv3* (magenta) and *her6* (grey) in control (top) and MBSm (bottom) embryos at 28hpf; transversal view; scale bar 30µm; annotations denote dorsal (D) and ventral (V). (a.ii) Magnification of inset from merged *gfap/elavl3* in **(a.i)** showing CTRL (top panel) and MBSm (bottom panel); annotations represent neural progenitor zone (NP=*gfap(+)/elavl(-))*, transition zone (T=*gfap(+)/elavl3(+)*). **(a.iii)** Normalized intensity mean of *elavl3* and *gfap* distributed across dorsal to ventral axis shows similar intensity profile in CTRL (top) and MBSm (bottom). **(b.i)** Quantitative analysis of *gfap* (green) and *elavl3* (magenta) pattern of expression across the dorsal to ventral axis of the Her6 domain observed in individual control versus MBSm embryos at 52hpf; data represents mean and SD of 4-10 slices per embryo; region of interest (ROI) delineates high *elavl3* vs *gfap in* CTRL (T and N zones) while in MBSm only the T zone is observed. **(b.ii)** Average *gfap* and *elavl3* intensity distribution across multiple embryos in the region of interest shown in **(b.i). (c)** Chromogenic WMISH of neuroD4 observed in multiple uninjected, CTRL and MBSm embryos at 52hpf; anterior to the left. **(d.i)** Single cell time series of Her6::Venus intensities observed between 34hpf and 47hpf in one control (Her::Venus) versus one MBSm embryo. **(d.ii)** Quantification of average Her6::Venus single cell intensities shown in **(d.i)**.

**Supplementary MovieS1**. Longitudinal view of Her6::Venus expression observed in the midbrain and hindbrain of Zebrafish embryo starting from 25hpf, scale bar 50 microns. Video data corresponds to stills included in Fig. 2a.i.

**Supplementary MovieS2**. Transversal view of Her6::Venus embryo showing two distinct expression domains in hindbrain (r6). Video includes a 3D rotated view at 30hpf followed by the expression pattern over development. Video data represents complete Z-stack view corresponding to stills in Fig. 2b.i 30-40hpf.

**Supplementary MovieS3.** Her6::Venus expression observed with single cell resolution in the hindbrain (r6) of embryo starting from 34hpf (corresponding to time point 0), scale bar 10 microns. Video data represents complete Z-stack corresponding to stills in Fig. 3c.

Fig. S1

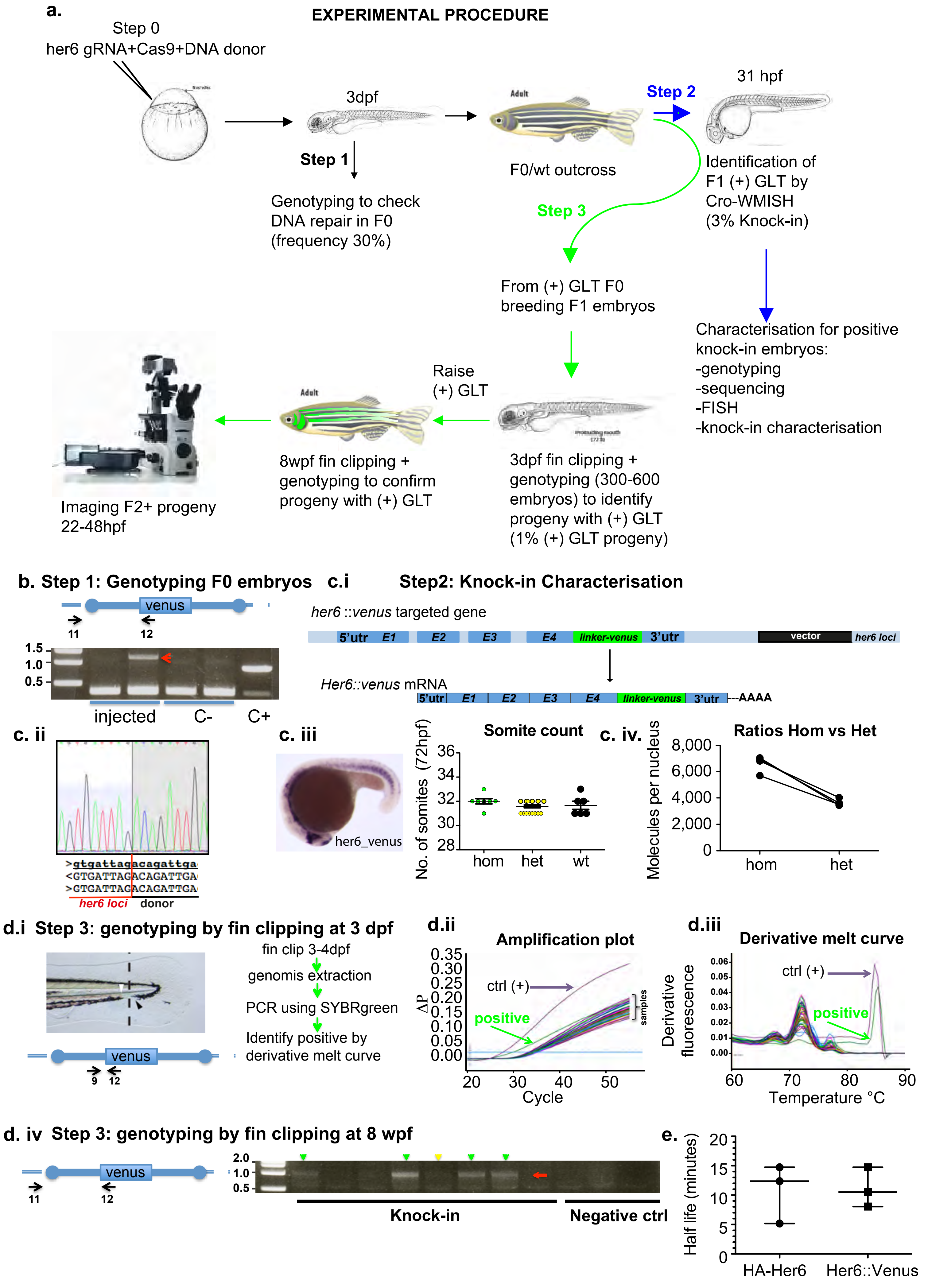
a.
EXPERIMENTAL PROCEDURE
Step 0
her6 gRNA+Cas9+DNA donor
3dpf
Step 1
Genotyping to check DNA repair in F0 (frequency 30%)
Adult
F0/wt outcross
Step 2
31 hpf
Identification of F1 (+) GLT by Cro-WMISH (3% Knock-in)
Step 3
From (+) GLT F0 breeding F1 embryos
Characterisation for positive knock-in embryos:
-genotyping
-sequencing
-FISH
-knock-in characterisation
Raise (+) GLT
Adult
Imaging F2+ progeny 22-48hpf
8wpf fin clipping + genotyping to confirm progeny with (+) GLT
3dpf fin clipping + genotyping (300-600 embryos) to identify progeny with (+) GLT (1% (+) GLT progeny)
b. Step 1: Genotyping F0 embryos
venus
11
12
1.5
1.0
0.5
injected
C-
C+
c.i
Step2: Knock-in Characterisation
her6 ::venus targeted gene
5'utr E1 E2 E3 E4 linker-venus 3'utr
vector
her6 loci
Her6::venus mRNA
5'utr E1 E2 E3 E4 linker-venus 3'utr ---AAAA
c. ii
>gtgattagacagattga
<GTGATTAGACAGATTGA
>GTGATTAGACAGATTGA
her6 loci
donor
c. iii
her6_venus
Somite count
No. of somites (72hpf)
36
34
32
30
28
hom
het
wt
c. iv.
Ratios Hom vs Het
Molecules per nucleus
8,000
6,000
4,000
2,000
0
hom
het
d.i Step 3: genotyping by fin clipping at 3 dpf
fin clip 3-4dpf
genomis extraction
PCR using SYBRgreen
Identify positive by derivative melt curve
venus
9
12
d.ii
Amplification plot
ΔP
0.35
0.30
0.25
0.20
0.15
0.10
0.05
0.00
ctrl (+)
positive
samples
20
30
40
50
Cycle
d.iii
Derivative melt curve
Derivative fluorescence
0.06
0.05
0.04
0.03
0.02
0.01
0.00
ctrl (+)
positive
60
70
80
90
Temperature °C
d. iv Step 3: genotyping by fin clipping at 8 wpf
venus
11
12
2.0
1.0
0.5
Knock-in
Negative ctrl
e.
Half life (minutes)
20
15
10
5
0
HA-Her6
Her6::Venus

Fig. S2

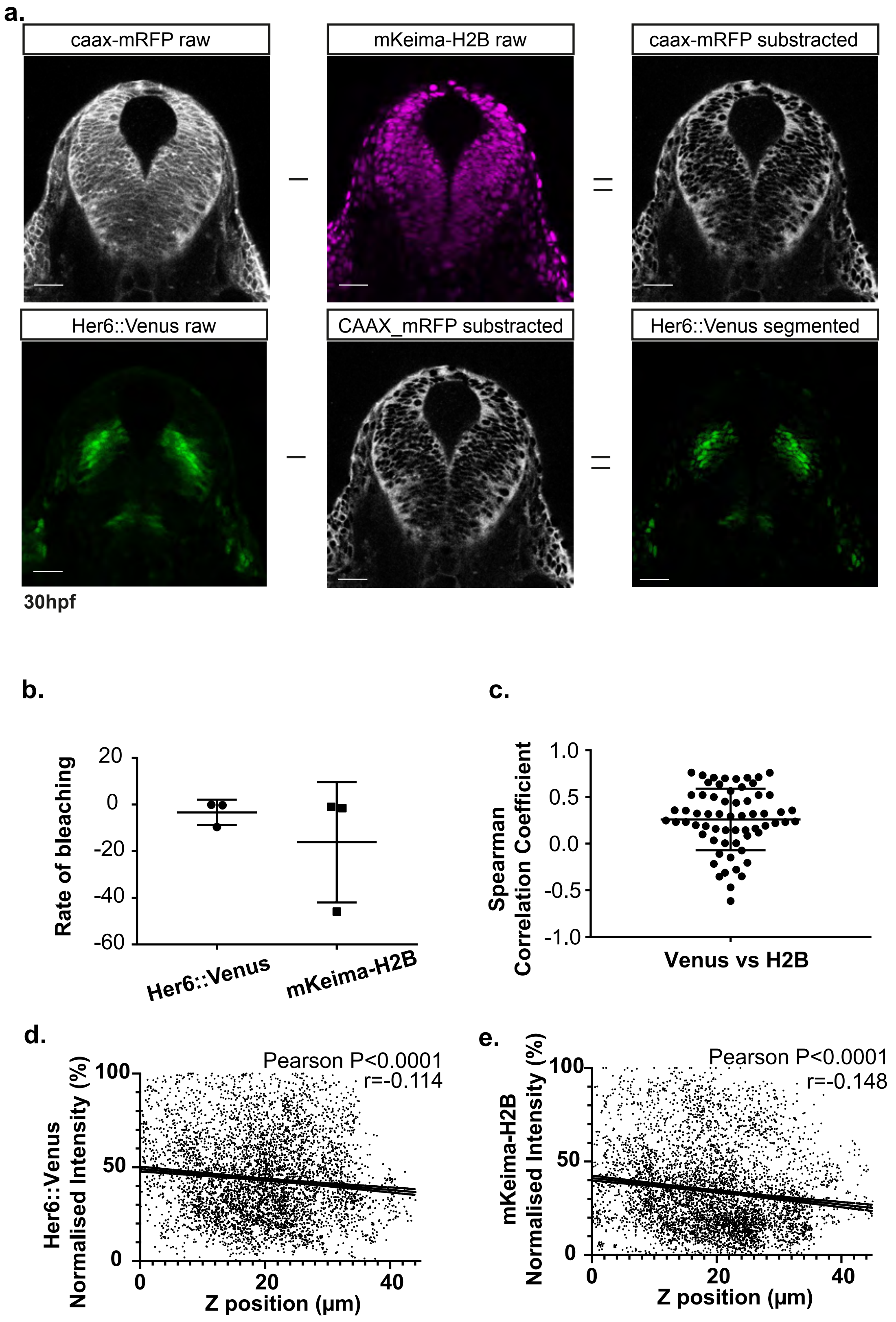
a.
caax-mRFP raw
mKeima-H2B raw
caax-mRFP substracted
Her6::Venus raw
CAAX_mRFP substracted
Her6::Venus segmented
30hpf
b.
Rate of bleaching
20
0
-20
-40
-60
Her6::Venus
mKeima-H2B
c.
Spearman Correlation Coefficient
1.0
0.5
0.0
-0.5
-1.0
Venus vs H2B
d.
Her6::Venus Normalised Intensity (%)
Pearson P<0.0001
r=-0.114
100
50
0
0
20
40
Z position (μm)
e.
mKeima-H2B Normalised Intensity (%)
Pearson P<0.0001
r=-0.148
100
50
0
0
20
40
Z position (μm)

Fig. S3

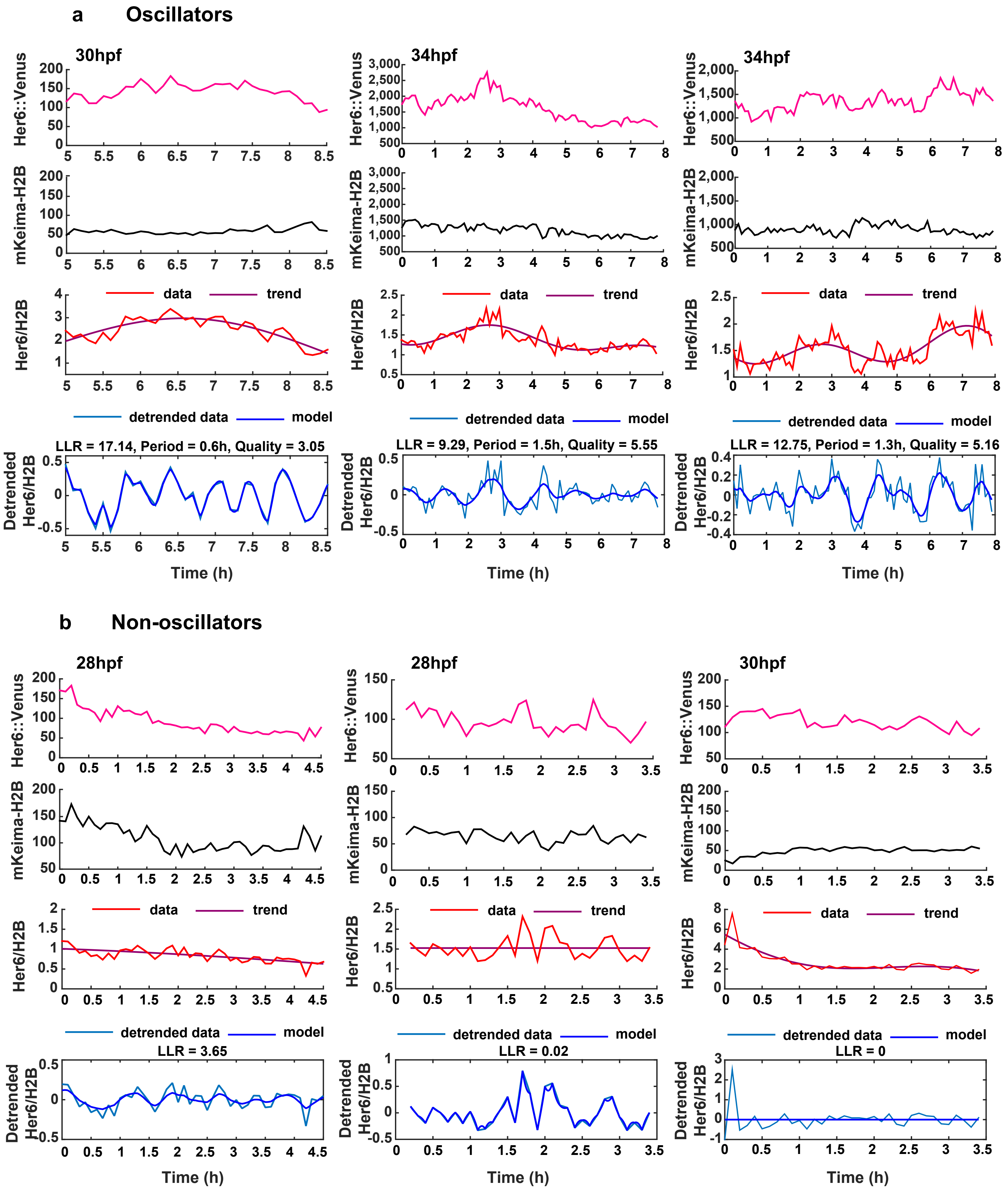
a
Oscillators
30hpf
34hpf
34hpf
Her6::Venus
mKeima-H2B
Her6/H2B
data
trend
detrended data
model
LLR = 17.14, Period = 0.6h, Quality = 3.05
LLR = 9.29, Period = 1.5h, Quality = 5.55
LLR = 12.75, Period = 1.3h, Quality = 5.16
Detrended Her6/H2B
Time (h)
b
Non-oscillators
28hpf
28hpf
30hpf
LLR = 3.65
LLR = 0.02
LLR = 0

Fig. S4

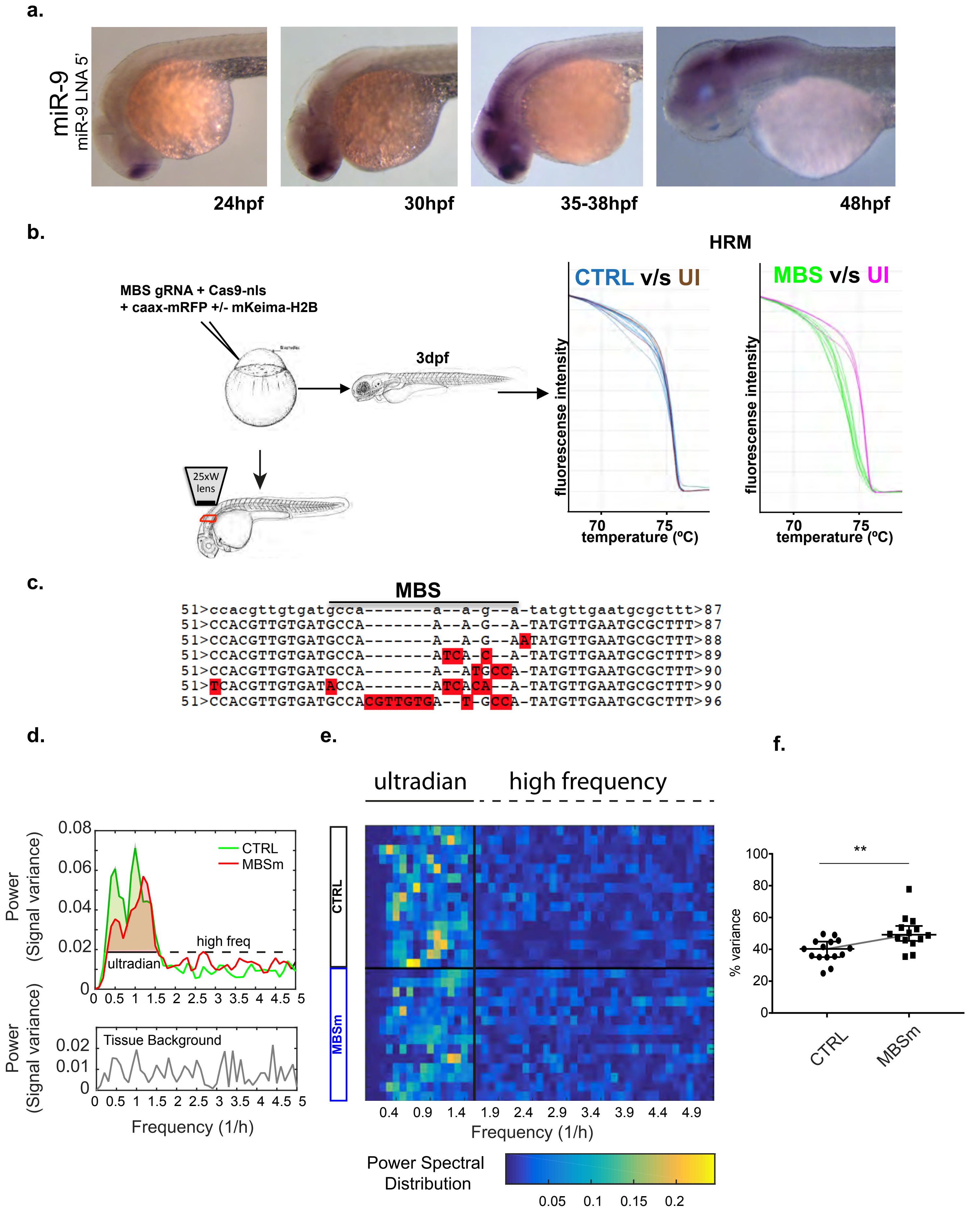
a.
miR-9
miR-9 LNA 5'
24hpf
30hpf
35-38hpf
48hpf
b.
MBS gRNA + Cas9-nls
+ caax-mRFP +/- mKeima-H2B
3dpf
25xW lens
HRM
CTRL v/s UI
MBS v/s UI
fluorescense intensity
70
75
temperature (ºC)
c.
MBS
51>ccacgttgtgatgcca-------a--a-g--a-tatgttgaatgcgcttt>87
51>CCACGTTGTGATGCCA-------A--A-G--A-TATGTTGAATGCGCTTT>87
51>CCACGTTGTGATGCCA-------A--A-G--AATATGTTGAATGCGCTTT>88
51>CCACGTTGTGATGCCA-------ATCA-C--A-TATGTTGAATGCGCTTT>89
51>CCACGTTGTGATGCCA-------A--ATGCCA-TATGTTGAATGCGCTTT>90
51>TCACGTTGTGATACCA-------ATCACA--A-TATGTTGAATGCGCTTT>90
51>CCACGTTGTGATGCCACGTTGTGA--T-GCCA-TATGTTGAATGCGCTTT>96
d.
Power (Signal variance)
0.08
0.06
0.04
0.02
0
CTRL
MBSm
high freq
ultradian
0 0.5 1 1.5 2 2.5 3 3.5 4 4.5 5
Tissue Background
0.02
0.01
0
Frequency (1/h)
e.
ultradian
high frequency
CTRL
MBSm
0.4 0.9 1.4 1.9 2.4 2.9 3.4 3.9 4.4 4.9
Frequency (1/h)
Power Spectral Distribution
0.05 0.1 0.15 0.2
f.
**
% variance
100
80
60
40
20
0
CTRL
MBSm

### Fig. S5

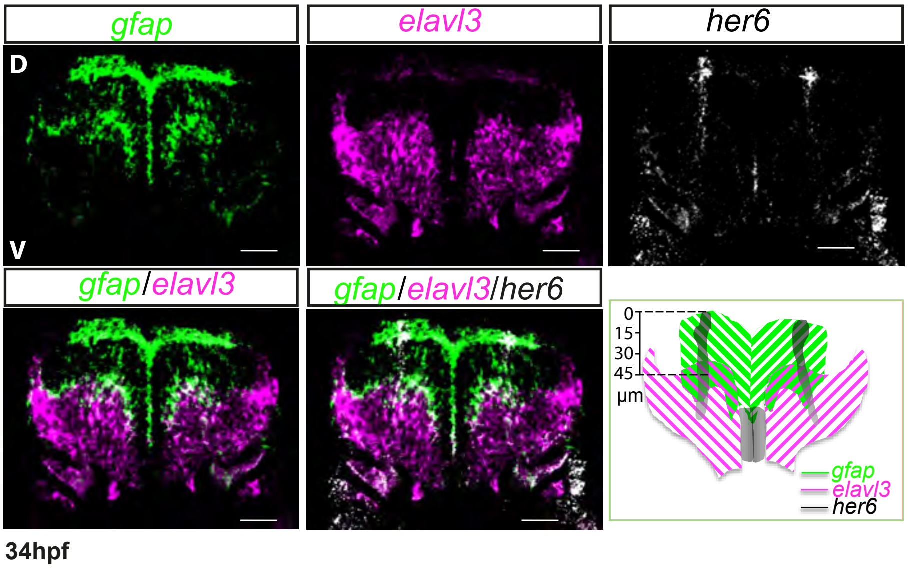
gfap
elavl3
her6
D
V
gfap/elavl3
gfap/elavl3/her6
0
15
30
45
µm
gfap
elavl3
her6
34hpf

Fig. S6

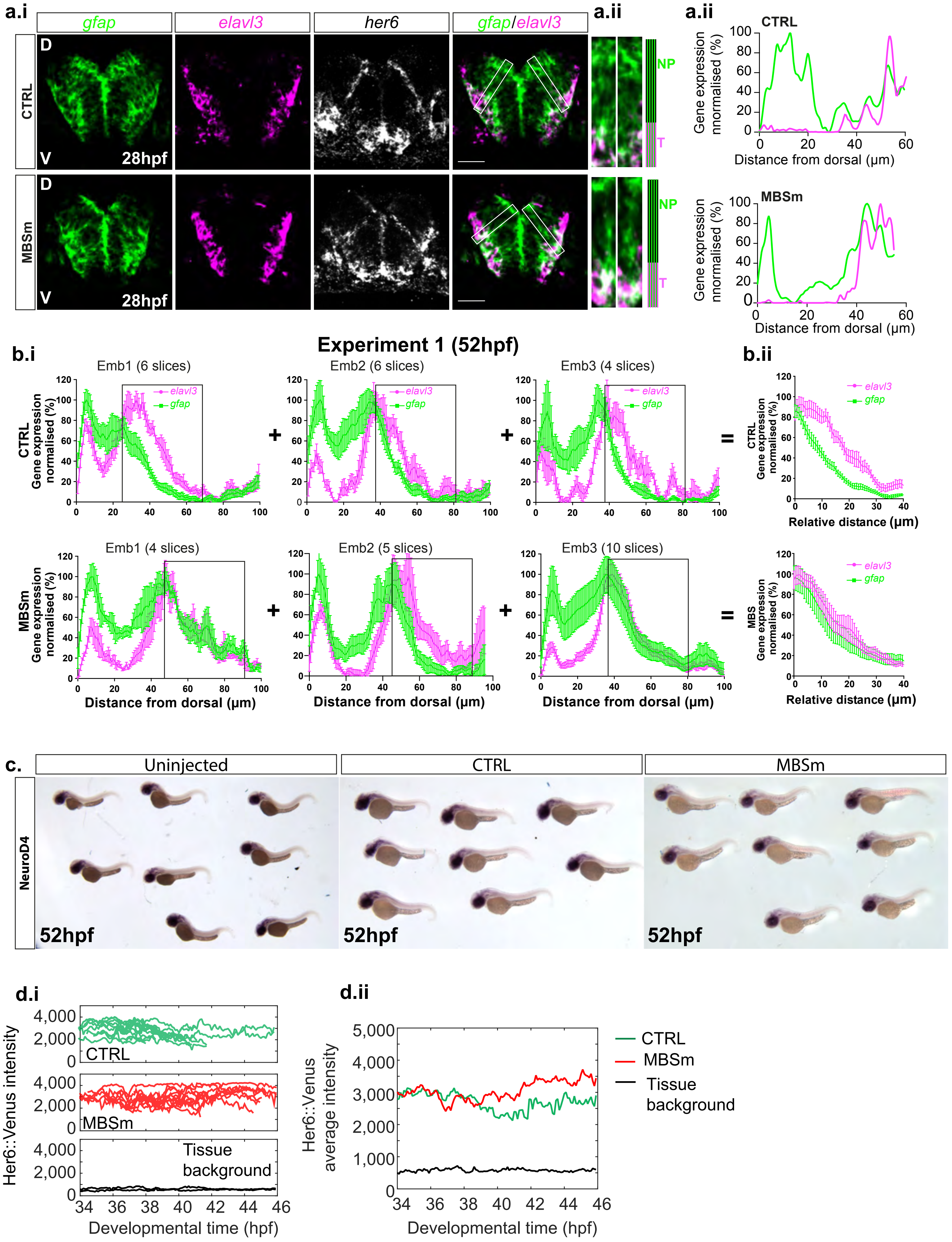

a.i
gfap
elavl3
her6
gfap/elavl3
CTRL
MBSm
D
V
28hpf
a.ii
NP
T
CTRL
MBSm
Gene expression nnormalised (%)
Distance from dorsal (μm)
b.i
Experiment 1 (52hpf)
Emb1 (6 slices)
Emb2 (6 slices)
Emb3 (4 slices)
Emb1 (4 slices)
Emb2 (5 slices)
Emb3 (10 slices)
CTRL Gene expressed normalised (%)
MBSm Gene expressed normalised (%)
Distance from dorsal (μm)
b.ii
Relative distance (μm)
c.
Uninjected
CTRL
MBSm
NeuroD4
52hpf
d.i
Her6::Venus intensity
Tissue background
Developmental time (hpf)
d.ii
Her6::Venus average intensity
CTRL
MBSm
Tissue background
Developmental time (hpf)
